# The mouse cortico-tectal projectome

**DOI:** 10.1101/2020.03.24.006775

**Authors:** Nora L. Benavidez, Michael S. Bienkowski, Neda Khanjani, Ian Bowman, Marina Fayzullina, Luis Garcia, Lei Gao, Laura Korobkova, Lin Gou, Kaelan Cotter, Marlene Becerra, Sarvia Aquino, Chunru Cao, Nicholas N. Foster, Monica Y. Song, Bin Zhang, Seita Yamashita, Muye Zhu, Darrick Lo, Tyler Boesen, Brian Zingg, Anthony Santarelli, Ian R. Wickersham, Giorgio A. Ascoli, Houri Hintiryan, Hong-Wei Dong

## Abstract

The superior colliculus (SC) is a midbrain structure that receives diverse and robust cortical inputs to drive a range of cognitive and sensorimotor behaviors. However, it remains unclear how descending cortical inputs arising from higher-order associative areas coordinate with SC sensorimotor networks to influence its outputs. In this study, we constructed a comprehensive map of all cortico-tectal projections and identified four collicular zones with differential cortical inputs: medial (SC.m), centromedial (SC.cm), centrolateral (SC.cl) and lateral (SC.l). Computational analyses revealed that cortico-tectal projections are organized as multiple subnetworks that are consistent with previously identified cortico-cortical and cortico-striatal subnetworks. Furthermore, we delineated the brain-wide input/output organization of each collicular zone and described a subset of their constituent neuronal cell types based on distinct connectional and morphological features. Altogether, this work provides a novel structural foundation for the integrative role of the SC in controlling cognition, orientation, and other sensorimotor behaviors.

## INTRODUCTION

Cortico-tectal projections from the cerebral cortex to the superior colliculus (SC) relay information to mediate a range of multimodal and cognitive functions^1–3^. In order for the SC to coordinate complex animal behaviors, such as attention, navigation, defense and decision-making, it requires the alignment and integration of top-down higher-order cortical information with sensorimotor maps. Disruptions in cortico-tectal projections have been implicated in attention-deficit and autism spectrum disorders^4–8^. Topographic cortico-tectal projections from primary visual, auditory, and motor areas have been well studied; however, the integration of higher-order associative cortical inputs to the SC has not been thoroughly characterized^3,9,10^. The mouse SC is an emergent structural model for the rodent visual system and the circuit formation that facilitates multisensory integration^11^; thus, it is important to understand the organizational principles of how the confluence of higher-order cortical information is integrated and channeled to downstream motor systems to regulate a diverse repertoire of behaviors.

It is well appreciated from early anatomical and electrophysiological studies in monkeys, cats, and other mammalian and non-mammalian species that the SC plays a prominent role in visuomotor and orienting behaviors^12,13^. With the prolific rise in functional studies on rodents, connectivity of the SC is often reviewed and generalized across multiple species to suggest there is little left to understand about its mesoscale connections^11^. However, to date, there is no complete anatomical map of SC connectivity in mice or any other species^14,15^. Current advancements in mapping the mouse connectome using modern neuroanatomic and computational tools facilitate a high throughput approach to assemble the first detailed connectivity map of descending cortical subnetworks to the SC: the cortico-tectal projectome. This provides a foundational structural model for the mouse SC that supports working hypotheses of regionally defined functions (such as flight vs. freezing or approach vs. aggression), and network- and cell type-specific functions. As such, additional efforts are still required to attain a comprehensive understanding of brain-wide subnetworks and their convergent and divergent information processing onto the complex SC laminar architecture.

The SC is composed of seven alternating fibrous and cellular layers, including three superficial (zo, zonal; sg, superficial grey; op, optic), two intermediate (ig, intermediate grey; iw, intermediate white), and two deep layers (dg, deep grey; dw, deep white)^16^. Functional studies often bisect the SC into medial and lateral halves; however, these broad undefined divisions do not sufficiently reflect how distinct subnetworks of contiguous or segregated inputs/outputs are distributed throughout the entire tectal volume. Thus, to systematically map the cortico-tectal projectome and define SC subregions, we characterized layer-specific cortical fiber terminations onto the entire SC. Computational analysis of the distribution of cortico-tectal fibers revealed that the SC can be subdivided into at least four radially distinct zones that extend along the medial-lateral and rostral-caudal axes, where each zone is defined by a unique subnetwork of cortical inputs. For each newly defined SC zone, we describe the complex alignment of sensory, somatic sensorimotor, and higher-order cortico-tectal projections that coincide with functional subnetworks for navigation, defense, decision-making, and action selection^1,17,18^.

## RESULTS

### General strategy for delineating the cortico-tectal projectome

First, we systematically analyzed and annotated the projection patterns of 86 cortico-tectal pathways from 150 from injections placed across the entire neocortex in adult mice (**Supplementary Figure 1**). Single anterograde tracer injections of either *Phaseolus vulgaris* leucoagglutinin (PHAL), or adeno-associated viruses expressing green (AAV-GFP) or red (AAV-tdTomato) fluorescent protein were confined to a single delineated cortical structure to produce region-specific projection terminal patterns in the SC (**Figure 1**). We also carried out double or triple tracer injections with combinations of PHAL, AAV-GFP and AAV-tdTomato to determine direct spatial correlations of axonal terminals arising from different cortical areas. Anterograde injections in the gustatory (GU), visceral (VISC), entorhinal (ENT), perirhinal (PERI), ectorhinal (ECT) areas, and claustrum (CLA) did not produce any evident projections to SC. The complete list of abbreviations and nomenclature can be found in **Supplementary Table 1**. All cases collected throughout this study, including injection sites and tracer details, can be found in **Supplementary Table 2.**

**Figure 1.**
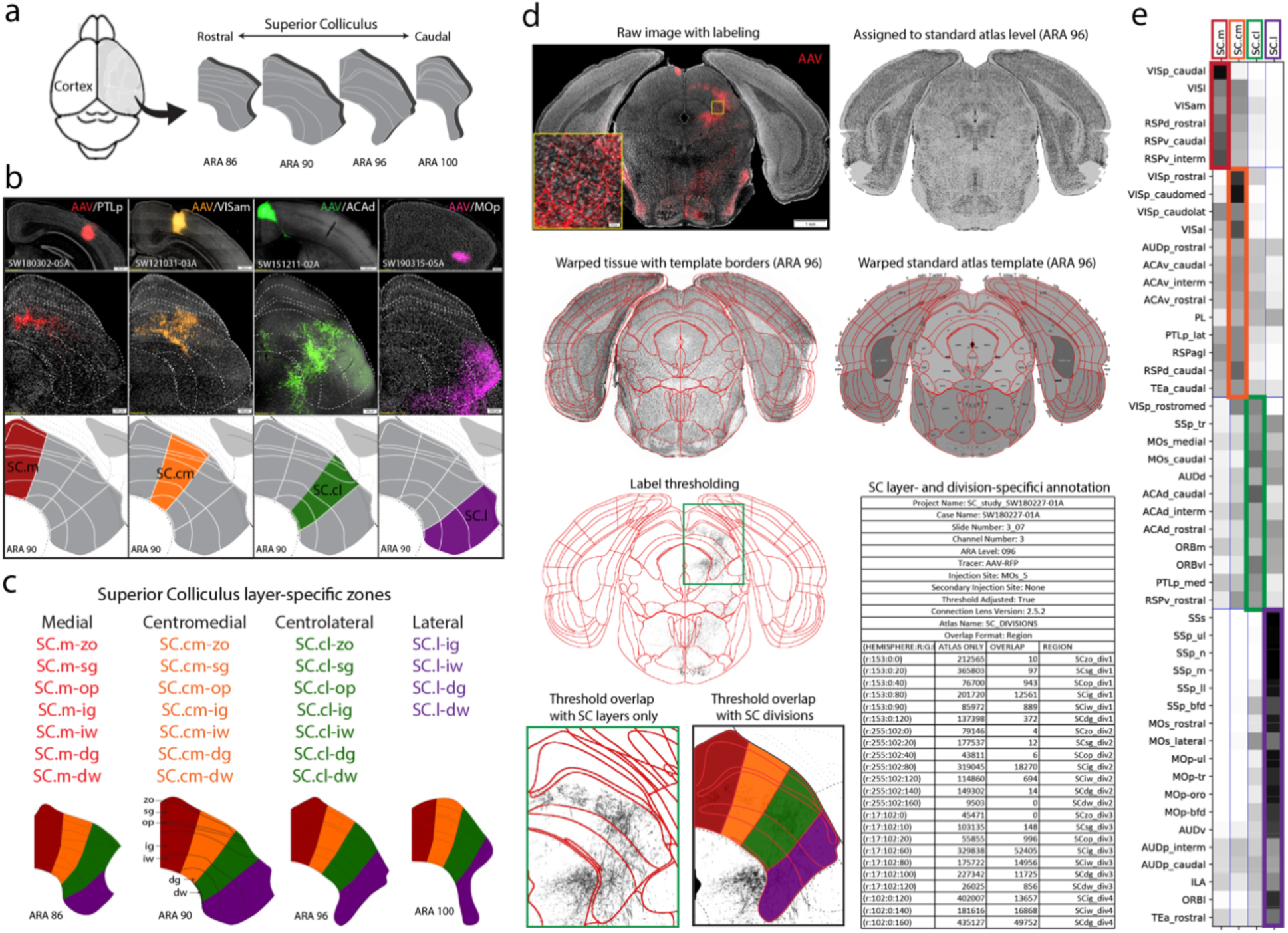
Experimental Workflow. **a)** Schematic of dorsal view of mouse brain cortex with arrow to map cortical projections across rostral to caudal levels of SC. **b)** Raw data examples of cortico-tectal projections targeting distinct zones within SC layers. Anterograde AAV injections into PTLp (red), VISam (orange), ACAd (green) and MOp (purple) terminate with different laminar and regional patterns in SC (ARA 90). Together, these representative cortico-tectal projections reveal layer-specific terminals distributed across four distinct zones along the medial-lateral axis of SC. **c)** Layer- and zone-specific SC nomenclature to facilitate referencing and quantification. **d)** The neuroinformatics workflow: Raw tissue image with anterograde labeling is assigned to the corresponding standard atlas level (this example is ARA 96). The tissue is warped based on template atlas borders and reconstructed using in-house neuroinformatics software, Connection Lens. Thresholded images are overlapped onto a custom atlas for zone- and layer-specific registration for pixel quantification. **e)** Re-ordered weighted matrix shows cortical ROIs along the y axis and the four SC zones along the x axis. Colored boxes along diagonal represent distinct cortico-tectal subnetworks. *Abbreviations: ARA, Allen reference atlas; AAV, Adeno-associated virus; PHAL, Phaseolus vulgaris leucoagglutinin. SC Layers: zo, zonal; sg, superficial grey; op, optic; ig, intermediate grey; iw, intermediate white; dg, deep grey; dw, deep white.*

Raw microscopy images with tracer-labeled cortico-tectal projections were assigned to ten corresponding adult mouse brain Allen Reference Atlas (ARA) templates (levels 84-102)^19^. Based on our initial observations of the cortico-tectal distribution patterns, we hypothesized that four distinct regional connectivity zones exist along the medial to lateral axis that follow dorsal to ventral radial boundaries (**Figure 1b**). Each zone resembles a radial column spanning the dorsal superficial layers to ventral deep layers: medial (SC.m), centromedial (SC.cm), centrolateral (SC.cl) and lateral (SC.l). We delineated the four zone boundaries onto ARA levels 86, 90, 96 and 100 to create a custom SC atlas, and designated zone-specific layer nomenclature to facilitate more detailed referencing and quantification cortical inputs of SC (**Figure 1c**).

To test that the four delineated SC zones contained distinct cortico-tectal inputs, we applied our Connection Lens informatics workflow to analyze 86 representative cortico-tectal pathways that cover the rostral-to-caudal and medial-to-lateral extent of the cerebral cortex and SC (**Supplementary Figure 1**). The selected images for analysis were registered to their corresponding ARA level and custom SC atlas template. Anterograde labels were then thresholded, graphically reconstructed, and overlapped onto the layer- and zone-specific custom SC atlas template. Cortico-tectal projection distributions and density for each experimental case were quantified based on the thresholded pixels overlapped within each SC zone and layer (**Figure 1d; Methods**). Quantified pixel densities across all cases were used as input values in the construction of weighted connectivity matrices to visualize all cortico-tectal inputs across distinct zones. An implementation of the Louvain algorithm was applied to the connectivity matrix for modular community detection of cortical injection sites with terminal distribution to common SC zones^20,21^ (**Figure 1e**). This analysis confirmed that our hypothesized custom SC atlas boundaries reflect unique topographic alignments of integrative cortical communities composed of higher-order associative cortical areas with visual, somatosensory and motor areas. Experimental data used for these analyses are openly available online at www.MouseConnectome.org.

Next, we analyzed whether brain-wide connectivity with SC zones would corroborate these cortico-tectal pathways. We targeted anterograde and retrograde injections in each SC zone and into thalamus, substantia nigra, zona incerta, and other hindbrain areas to map their topographic patterns. Finally, we reconstructed thalamic projecting SC neurons to characterize their morphometric profiles and to assess whether individual neurons were segregated by discrete zones. All neuronal morphologies are cataloged into a digital library, accessible on www.NeuroMorpho.Org^22^.

### Sensory, somatosensory, and motor projections to SC

The network organization of cortico-tectal projections reflect the topographic representations of visual, auditory, somatosensory and motor maps conserved across both neocortex and SC. To better understand how higher-order inputs integrate with multisensory information within our new SC zones, we first established and quantified a distribution map for each sensory modality (**Figure 2**). We found that cortico-tectal subnetworks have similar organization principles as those identified in the cortico-cortical and cortico-striatal projectome^23,24^. The input patterns support the specialized mechanisms needed for laminar and regional integration within the SC.

**Figure 2.**
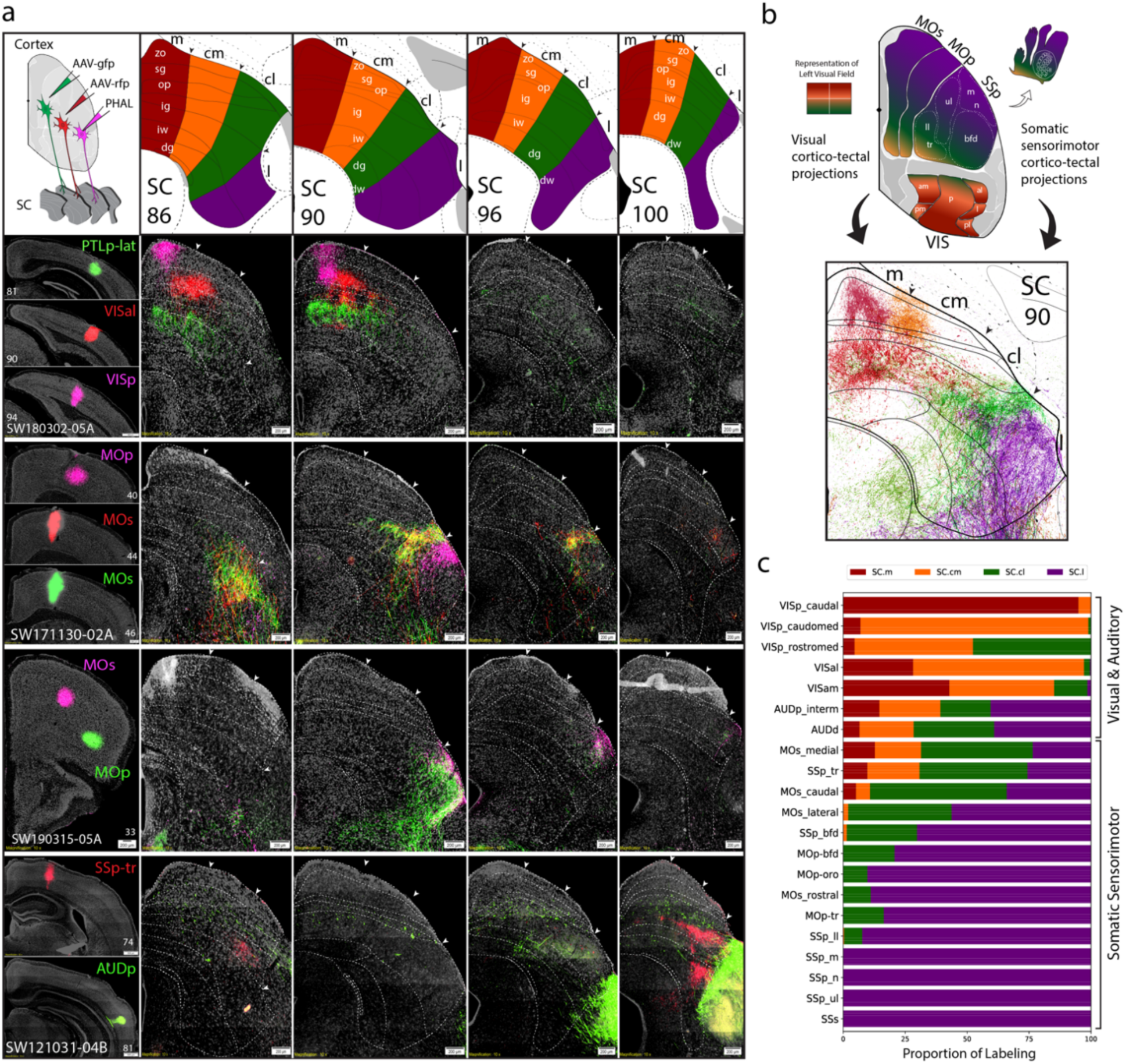
Visual, auditory, and somatic sensorimotor cortical projections to SC zones. **a)** Schematic of injection strategy using triple or double anterograde tracers into cortex. Top row: VISp, VISal, and PTLp-lat projections to SC.m and SC.cm. Second row: two neighboring caudal MOs regions and a rostral MOp injection send projections to adjacent, but non-overlapping zones in the SC.cl and SC.l. Third row: rostral MOs and rostral-ventral MOp projections to the SC.l zone. Bottom row: SSp-tr and AUDp projections target caudal SC.l zones. **b)** Color-coded schematic overview of the left visual field and body part topography based on their target zones in SC at level 90. Reconstructions below. **c)** Stacked bar chart for visualization of proportion of labeling across each SC zone (*x* axis) from each cortical ROI (*y* axis) for each selected ROI. Values represent proportion of pixel density for an individual cortical ROI (n=1 per row) across each SC zone. See **Supplementary Table 3** for zone- and layer-specific values. SC boundaries in all figures were delineated based on Nissl-stained cytoarchitecture. Dashed lines correspond to specific layers in each SC level.

As observed in previous studies, projections from the primary visual cortical area (VISp) to superficial SC layers align through intrinsic connectivity with intermediate layers which may coordinate a range of visually guided responses^25–28^. Consistent with previous studies, our results display that SC.m and SC.cm receive input from all visual cortices with a conserved representation of the upper central and upper peripheral visual field (**Figure 2a**). The VISp exclusively targets superficial layers, while the secondary visual areas, including the anterolateral (VISal), lateral (VISl), anteromedial (VISam), and posteromedial (VISpm), predominantly target the intermediate layers. SC.cl receives VISp inputs representing the lower central and lower peripheral field (**Figure 2b**). Notably, no VISp projections are found in SC.l.

Within the cortex, somatic sensorimotor areas are each organized into five distinct highly interconnected subnetworks that integrate the somatotopic body map and project subcortically to the caudoputamen (CP) along the dorsolateral and ventrolateral domains (CPi.dl and CPi.vl)^23,24^. Within each somatotopic subnetwork, primary somatosensory (SSp), supplementary somatosensory (SSs), primary motor (MOp) and secondary motor (MOs) areas send parallel descending projections to SC zones. The rostral SC.l dominantly receives inputs from SSp-m (mouth), SSp-n (nose), and SSp-ul (upper limb), with inputs to both SC.cl and SC.l from SSp-bfd (barrel field). The SSp-ll (lower limb) and SSp-tr (trunk) project to the caudal levels of SC.cl and SC.l (**Figure 2a**). Similarly, all MOp body regions predominantly project across the SC.l overlapping with their counterpart SSp inputs. Rostral-lateral MOs regions project to SC.cl and SC.l; and MOs-frontal eye field (MOs-fef)/ACAd sends tiered projections to the SC.cl (**Figure 2b**).

### Higher-order associative and prefrontal cortical projections to SC

Next, we mapped the projections of all higher-order associative cortical areas to SC to investigate how they integrated with multimodal and somatic sensorimotor maps^9^. These areas including, the anterior cingulate (ACA), retrosplenial (RSP), posterior parietal (PTLp), temporal association (TEa), and prefrontal (PFC) have preferential targets across SC.

Anterograde injections into the granular RSPd (dorsal) and RSPv (ventral) regions, and RSPagl (agranular) produce relatively confined distribution patterns within the SC zones (**Supplementary Figure 3a**). SC.m-sg/op/ig and SC.cm-sg/op/ig predominantly receive inputs from RSPagl and RSPd. These RSP→SC.m/cm projections align with the upper peripheral and central visual fields from VISp (**Figure 3a**). SC.cl receives more terminals from RSPv than RSPd and RSPagl.

**Figure 3.**
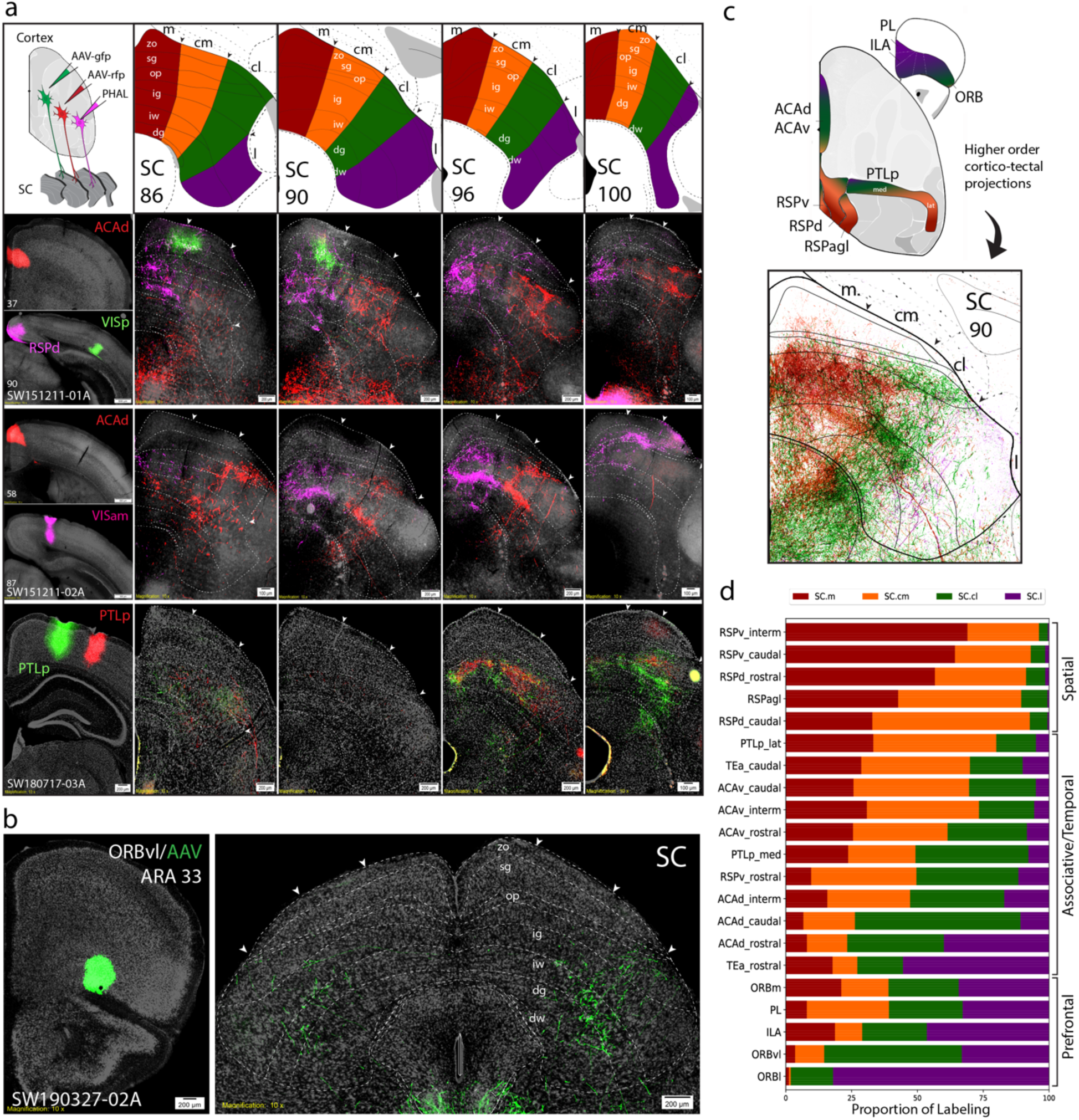
Distribution of higher-order cortical inputs across SC zones. **a)** Top row: VISp, RSPd and ACAd-rostral projections to SC. Middle row: ACAd-interm and VISam projections. Bottom row: two PTLp-medial projections. **b)** ORBvl sends bilateral projections that ascend into SC.cl and SC.l. **c)** Schematic of higher-order association areas part of medial cortico-cortical subnetworks. Color-coding represents topography based on cortico-tectal projection patterns that target distinct SC zones. Reconstruction is composite of inputs to SC ARA 90. **d)** Stacked bar chart for visualization of proportion of labeling across each SC zone (*x* axis) from each cortical ROI (*y* axis) for each selected ROI. Values represent proportion of pixel density for an individual cortical ROI (n=1 per row) across each SC zone. See **Supplementary Table 4** for zone- and layer-specific values.

Anterograde tracer injections into the ACAv (ventral) and ACAd (dorsal) produce robust overlapping projections across SC with a tiered/columnar organization (**Figure 3a**). Topographic outputs from rostral to caudal ACAv project primarily to SC.cm and SC.m. ACAd-caudal projects most densely to SC.cl, while ACAd-rostral and intermediate project more uniformly across SC.cl and SC.l. Collectively, ACA processes attention-related and affective information associated with pain and aggressive behaviors, which is consistent with its medial to lateral integration across multimodal maps in SC^18^.

The PTLp is regionally subdivided into a medial/rostral part spanning ARA 75-80 (PTLp-med), and lateral/caudal part spanning ARA 81-94 (PTLp-lat). SC.m and SC.cm mainly receive inputs from PTLp-lat, which align more with other visual cortical inputs. SC.cl and SC.l mainly receive inputs from PTLp-med, which align with more motor and somatosensory inputs, including the ACAd (**Figure 3a,c,d**).

The SC also receives bilateral inputs from prefrontal areas (PFC), including orbitofrontal medial (ORBm), lateral (ORBl), ventrolateral (ORBvl), prelimbic (PL) and infralimbic (ILA) (**Figure 3b; Supplementary Figure 3b**). Our results reveal an ILA→SC.m/l pathway that is in register with the representation upper peripheral visual field and somatic sensorimotor subnetwork, suggesting a divergence of information across zones. From cortical areas along the lateral edge of neocortex, the temporal association area (TEa) sends distributed projections across each SC zone, with TEa-rostral clustered with caudal SC.l, and TEa-caudal projects to SC.cl (**Figure 3d**). Consistent with these results, at the cortical level, TEa-rostral also shares dense bi-directional connections with somatic sensorimotor areas; while TEa-caudal is heavily connected with the visual and auditory cortical areas^23^.

### Cortico-tectal projections organized as modular communities

To understand the modular subnetwork organization of the cortico-tectal system, we analyzed the distribution of cortico-tectal projections in the SC using the Louvain community detection algorithm (**Figure 4a,b**). The Louvain analysis revealed that the SC zones also constitute four modular subnetworks differentiated by cortico-tectal inputs to each of the SC rostrocaudal levels. SC.m and SC.l modular networks are highly differentiated, whereas SC.cm and SC.cl are distinct, but closely related.

**Figure 4.**
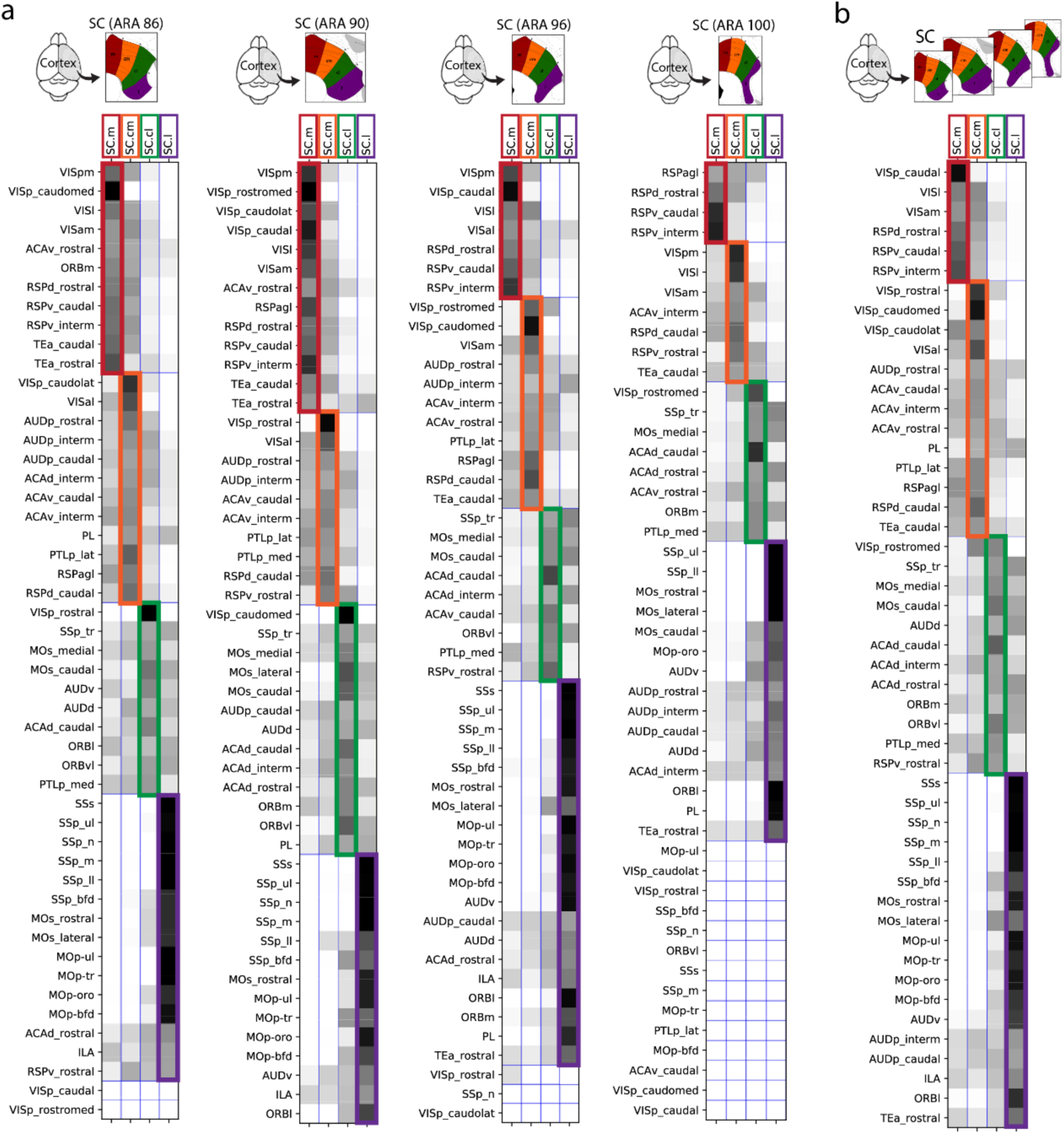
Visualization of cortico-tectal modular communities at each custom SC level. Matrices show communities identified by Louvain analysis of cortical injection ROIs (*y* axis) and SC zones (*x* axis). Edges are shaded according to their connectivity labeling, and colored boxes along the diagonal reflect modular communities identified. Matrix community color scheme corresponds to SC custom atlas zones, and same colors are used across levels to compare community structures. **a)** Communities detected within each custom SC atlas level. Matrices with empty cells in the bottom do not contain labeling and thus are not clustered into communities. **b)** Combined matrix of consensus communities from all four levels.

The SC.m communities contained primarily inputs from visuospatial cortical areas, including primary and secondary VIS areas and RSP. By contrast, the SC.l communities contained the majority of inputs from somatic sensorimotor/prefrontal subnetworks from SSp, SSs, MOs, MOp, and PFC areas. This separation of communities suggests functionally distinct processing of visual information in the SC.m and sensorimotor information in the SC.l. In contrast, the SC.cl and SC.cm receive more integrated functional inputs. The SC.cl zone is aligned with the lower central visual field, and communities in the SC.cl contained visuospatial inputs from secondary motor area MOs-fef/ACAd, VISp-, and RSPv-rostral regions. In general, SC.cl receives motor inputs similar to SC.l, but also from other auditory and associative regions that are characteristic of SC.cm. The SC.cm communities contained multimodal and associative regions, but not prominent somatic sensorimotor cortical areas.

These results are complementary to our previous work which showed that a number of higher-order associative areas along the medial bank of the neocortex including, ACA, RSP, and PTLp, are organized into highly-connected medial cortico-cortical subnetworks that integrate and relay visual, auditory, somatosensory, and spatial information (from the subiculum via RSPv) to the PFC areas^23^. Similarly, outputs from the cortico-cortical somatic sensorimotor subnetwork converge predominantly into communities within the SC.l and SC.cl. Overall, our analysis indicates that cortico-tectal inputs conserve cortico-cortical subnetwork organization. Additional analysis from thalamic and brainstem injections corroborate these cortico-tectal Louvain communities in relation to brain-wide subnetworks.

### Brain-wide connectivity corroborates cortico-tectal networks

The SC relays integrated information as motor commands via direct projections to brainstem motor and thalamic nuclei^29,30^. Thus, after establishing the cortico-tectal projectome and identifying their convergence into distinct communities, we validated the input/output patterns for each SC zone. We investigated their correlation and brain-wide interactions with cortico-tecto-thalamic and cortico-tecto-basal ganglia subnetworks. Using anterograde and retrograde coinjections into the SC, we assembled a complete mesoscale connectivity diagram of the mouse SC that illustrates the inputs and outputs distributed across sensory, motor, and behavior state regions within the thalamus, hypothalamus, midbrain, pons, medulla and cerebellum (**Figure 5a; Figure 6**). We created an anterograde projection map from each SC zone which summarizes the spatial and topographic innervation patterns (**Figure 5b; Supplementary Figure 4-5**). This analysis reveals a systems level architecture of SC zones integrating higher-order associative and multimodal information throughout functionally implicated subnetworks.

**Figure 5.**
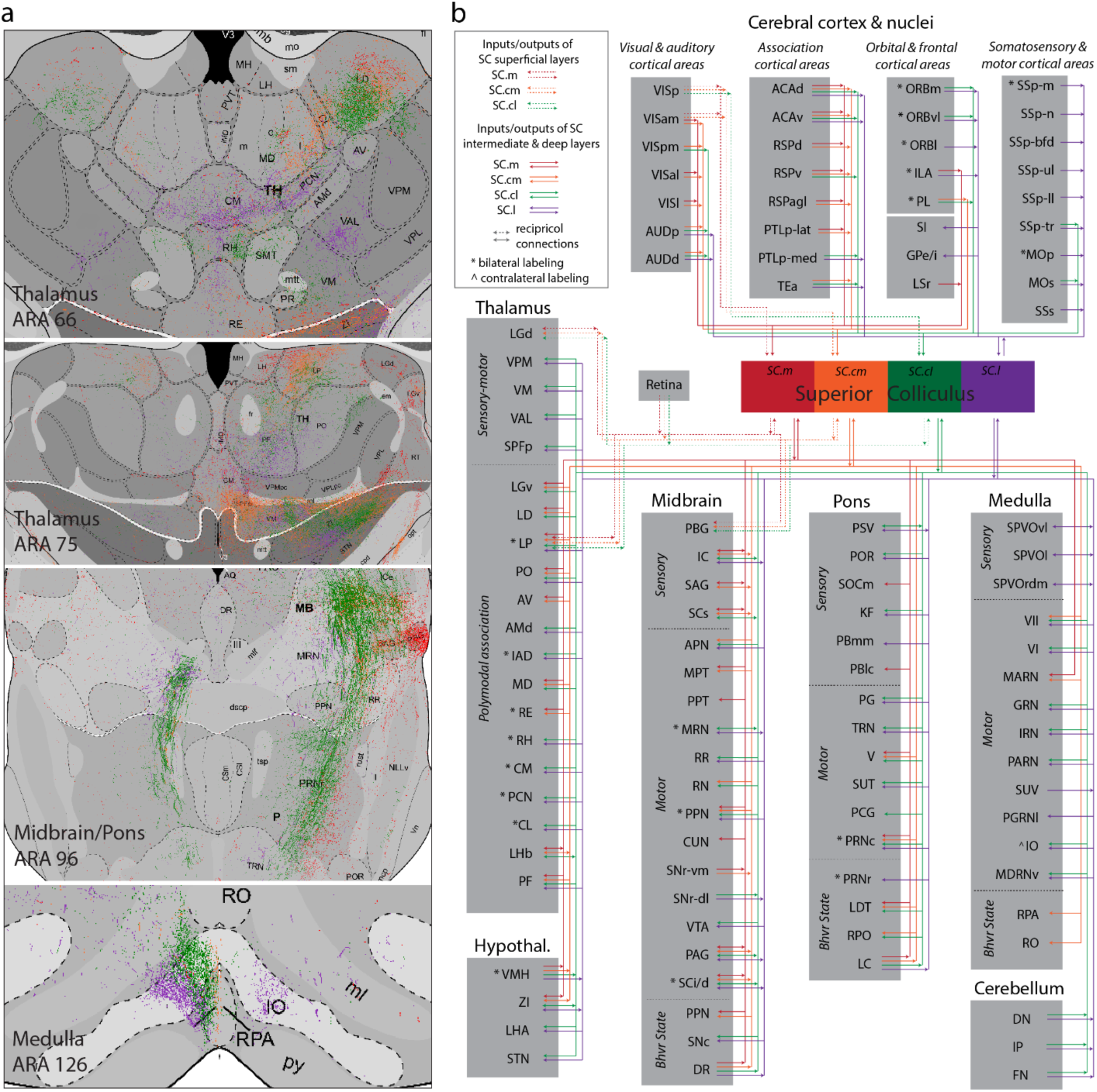
Connectome diagram of the mouse superior colliculus. **a)** Unweighted wiring map of all inputs and outputs of SC zones and layers. Assembled using data from anterograde and retrograde injections placed in SC to systematically trace outputs inputs distributed from cortex down through the hindbrain and cerebellar structures. Annotation table can be found in **Supplementary Table 5. b)** Color-coded map of anterograde projections from each SC zone across the brain demonstrating topographic output patterns.

**Figure 6.**
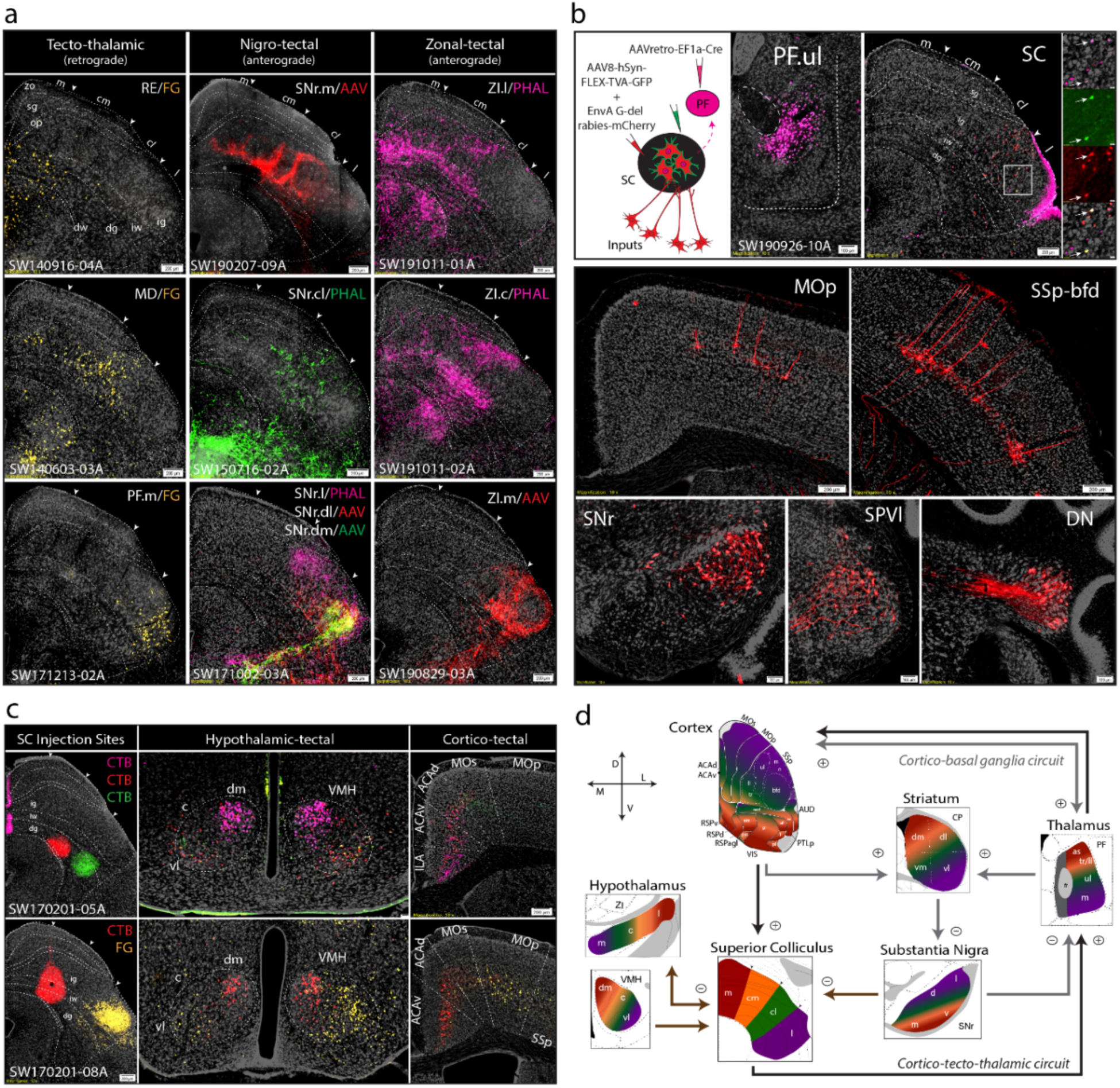
Brain-wide inputs and outputs of SC zones. **a)** Raw connectivity data in SC from nine separate anterograde or retrograde cases show topographic patterns across distinct SC zones from SNr, zona incerta (ZI) and thalamic nuclei (columns). Tecto-thalamic neurons were distributed heterogeneously across superficial, intermediate and deep layers, but clustered preferentially in distinct zones. Left column: retrogradely labeled cells from FG injections in different thalamic nuclei; RE projecting cells in SC.m/cm; MD projecting cells predominantly in SC.cl; and PF.m projecting cells from SC.l. Middle column: labeling produced from anterograde injections into distinct SNr domains project to SC. Right column: anterograde labeling from ZI.l targets SC.m/cm; ZI.c densely targets SC.cl; and ZI.m targets SC.l. **b)** TRIO-tracing strategy targeting AAV-retro-Cre in PF.ul (upper limb), and Cre-dependent rabies virus and helper virus in SC.l reveal mono-synaptic inputs to the PF-projecting neurons in the SC.l. **c)** Two separate cases with retrograde injections into SC from two separate cases. Cells are topographically labeled in VMH domains and cortex. **d)** Topographic organization of mouse SC connectivity. Directional cross at bottom left. The four zones of the SC are connected either uni- or bi-directionally with cortex, PF, ZI, VMH and SNr. See **Supplementary Table 1** for complete list of abbreviations.

#### Zone-specific cortico-tecto-thalamic subnetworks

We identified that each SC zone receives convergent inputs from a specific set of cortical areas as described above. Here, we demonstrate how each zone sends projections back to specific thalamic nuclei that send ascending projections to the same cortical areas, thus completing the loop within the cortico-tecto-thalamic subnetwork (**Figure 6**). The SC.m and SC.cm predominantly integrate VIS, RSP and PTLp-lat information through convergent inputs across layers. These cortico-tectal inputs (in SC.m/cm/cl) align with retrogradely labeled cells from separate FG injections into the reuniens (RE), mediodorsal (MD), laterodorsal (LD), and lateroposterior (LP) nuclei of the thalamus (**Figure 6a; Supplementary Figure 6a**). Labeling from RE was also traced back to prefrontal and associative cortical areas suggesting the RE may relay integrated visual information from the SC to these medial cortico-cortical subnetworks. This information ultimately sends feedback projections to SC (RSP/ACA/PFC→SC→RE→RSP/ACA/PFC), as part of the cortico-tecto-thalamic circuit (**Supplementary Figure 6d**).

Retrograde FG injections were targeted to somatic sensorimotor related thalamic nuclei including, ventrolateral part of parafascicular nucleus (PF.vl), the caudal ventromedial nucleus (VM), and paracentral nucleus (PCN) (**Figure 6a**). Cells were backlabeled in the SC.cl/l, as well as SSp-m, SSp-ul, and MOp-orofacial areas (**Supplementary Figure 6a**), which are all areas part of somatic sensorimotor subnetworks processing orofacial and upper-limb information. The caudal levels of SC.cl and SC.l (receiving more trunk and lower-limb information) send selective projections to the dorsal PF and rostral VM, suggesting cortico-tectal and tecto-thalamic pathways conserve topographic organization. Meanwhile, the corresponding thalamic nuclei and cortical areas also generate parallel mono- or multi-synaptic projections to functionally correlated subdomains of the basal ganglia, including the CP and SNr^23,24^.

#### Interactions between cortico-tectal and basal ganglia subnetworks

Next, we investigated the interactions of cortico-tectal and basal ganglia subnetworks. Each of the SC zones harboring distinct thalamic projecting neurons receives convergent inputs directly from functionally correlated cortical areas (as aforementioned) and domain-specific regions from SNr, that is, the motor output node of the basal ganglia network (**Supplementary Figure 6d**). Anterograde injections into ventromedial SNr send descending projections to SC.m, SC.cm, and SC.cl that align with patterns from cortico-tectal projections from the ACAd and RSPv. Anterograde injections into dorsolateral and dorsomedial SNr domains send descending inputs confined to SC.l that align with inputs from somatic sensorimotor cortical areas and PF-projecting SC neurons (**Figure 6a**).

To show the subnetwork interactions of this cortico-tecto-thalamic pathway and reveal mono-synaptic inputs to the PF-projecting neurons in the SC.l, we used a TRIO-tracing strategy targeting AAVretro-Cre in dorsomedial PF, and Cre-dependent rabies virus and helper virus in SC.l^31^ (**Figure 6b**). As expected, rabies labeled cells that target PF-projecting SC.l neurons were found in SSp-ul, SSp-m, SSs, and MOp-oro cortical areas. Additionally, neurons were labeled in the dorsolateral SNr, deep cerebellar nuclei, and the lateral part of the spinal motor nucleus of the trigeminal (SPV) associated with processing orofacial and tactile information^32^. Using this strategy, we further validated the multi-synaptic somatic sensorimotor cortico-tecto-thalamic subnetwork, and identified how cortical inputs are integrated with other information in the thalamic projecting SC neurons. These data provide evidence that suggests the SC.l receives convergent inputs from functionally correlated cortical areas, subsets of cortico-basal ganglia (i.e. dorsolateral SNr), and other subcortical structures associated with hand-to-mouth coordination behaviors.

To assess this circuit with an alternative approach, we injected two anterograde tracers (AAV-mCherry or PHAL) into the ORB and SNr, respectively, and a retrograde tracer (Rabies-GFP) into the MD (**Supplementary Figure 6b**). We found that in the SC, both cortical and nigral projection terminals directly innervate MD-projecting neurons in the SC.m and SC.cm, thereby supporting the specificity of this multi-synaptic cortico-tecto-thalamic pathway. This suggests that excitatory cortical inputs and inhibitory nigral inputs converge onto the same SC projection neurons, though further studies are needed.

#### Hypothalamic interactions with SC

The SC zones also receive topographic inputs from hypothalamic structures, including zona incerta (ZI), and ventromedial nucleus of hypothalamus (VMH)^33,34^. The output patterns of ZI to SC support the topographic organization of medial visuomotor and lateral somatic sensorimotor subnetworks. Anterograde injections into lateral ZI project to SC.m and SC.cm, central ZI projects to SC.cm and SC.cl, and medial ZI projects to SC.l (**Figure 6a**). Interestingly, lateral ZI→SC.m/cm innervate intermediate and superficial layers, further supporting the intrinsic alignment of visuomotor subnetworks that also align with our identified cortico-tectal communities.

The SC.cl and SC.l intermediate and deep layers receive dense inputs from the ventrolateral part of the ventromedial hypothalamus (VMH.vl) known to be involved in approach, appetitive, reproductive and social attack responses^35,36^. Triple retrograde injections in SC.m-ig (CTb-647), SC.cl-dg (CTb-555), and SC.l-dg (CTb-488) reveal that SC.m receives dense inputs primarily from VMH.dm, whereas SC.cl and SC.l receive more inputs from VMH.vl. Another experiment comparing SC.cm (CTb-555) and SC.l-ig (FG) revealed SC.cm receives input from VMH.dm/c, and SC.l receives input from VMH.vl. (**Figure 6c**). These findings are congruent with functional networks implicating the lateral SC in appetitive and approach behaviors^17,18^.

### Cataloging SC neuronal morphology

Finally, we characterized SC cell types based on their anatomical locations, projection targets, and neuronal morphology. Based on our findings from FG injections in thalamic nuclei retrogradely labeling cells in distinct SC zones (**Figure 6a; Supplementary Figure 6a**), we hypothesized that projection neurons located in the four SC zones were morphologically distinct. To test this, representative neurons in each zone were labeled via a G-deleted-rabies injection in the LD (for SC.m/cm), in the RE (for SC.m/cm/cl), and dorsal PF (intended for SC.cm/cl/l) (**Figure 7a**). The large injection into the dorsal PF also partially infected neurons in the midbrain pretectal region (MPT), which retrogradely labeled cells in the superficial layers of SC.m. We thus refer to this injection as PF/MPT in the analysis and interpretation of the results. All neuronal morphologies reconstructed in this study along with detailed experimental metadata are cataloged into a digital library, accessible on www.NeuroMorpho.Org^22^.

**Figure 7.**
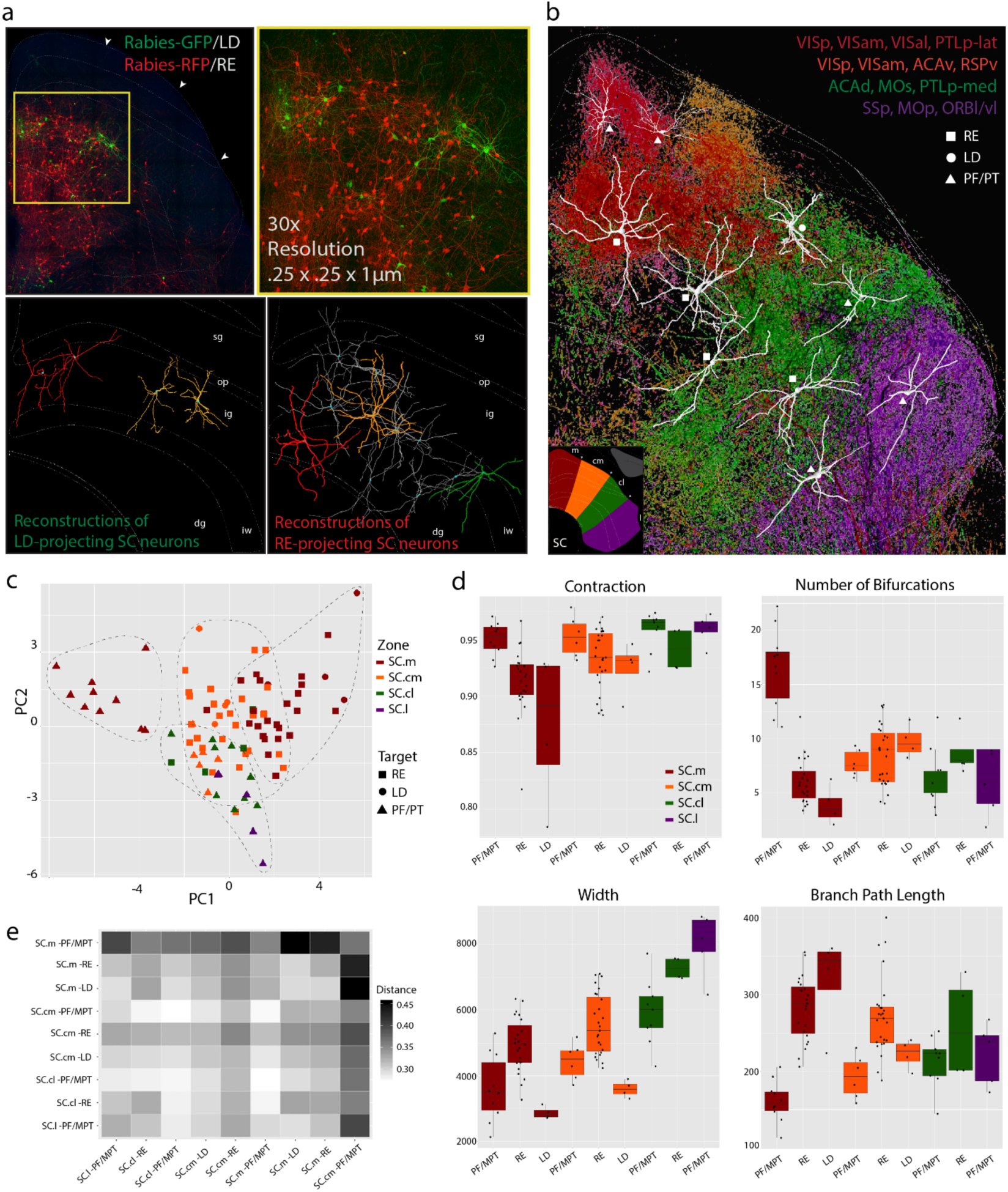
Characterization of neuronal cell types in the SC based on their anatomical locations, projection targets and neuronal morphology. **a)** LD projecting neurons in SC labeled with rabies-GFP in SC.m/cm-ig, and RE projecting neurons labeled with rabies-RFP injected into cells in SC.m/cm-ig/iw/dg. Tissue was processed by the SHIELD clearing protocol and followed by confocal imaging for 3D-reconstruction. Dendrites from LD and RE-projecting SC neurons were reconstructed to visualize differences and dendritic arborizations across SC layers and zones. **b)** Examples of reconstructed neurons were overlapped onto reconstructions of cortico-tectal projections on the SC atlas. This facilitates analysis of spatial correlation between cortical projection fields and “receptive fields” of SC-projection neurons. **c)** Principal component analysis (PCA) shows segregation of zone- and target-specific cells based on measured morphological features (see Methods). Number SC projection neuron reconstructions to RE (n=54), LD (n=8), and PF/MPT (n=31). **d)** Examples of reports from pairwise tests of morphological parameters that survived the false discovery rate correction tests. Significant group differences are presented with whisker plots, where the center line represents the median, box limits show the upper and lower quartiles and the whiskers represent the minimum and maximum values. P-values for all parameters can be found in **Supplementary Table 6. e)** Persistence-based neuronal feature vectorization framework was also applied to summarize differences between projection neurons from each zone to each thalamic region. The strength of differences is presented as a gradient with white showing no difference and black the greatest differences. *Abbreviations: LD, laterodorsal nucleus; PF/MPT, parafascicular nucleus of thalamus/midbrain pretectal area; RE, reuniens nucleus of thalamus.*

Reconstructions of rabies-labeled SC neurons projecting to RE, LD and PF/MPT allow us to analyze the spatial correlation between cortical projection fields and “receptive fields” of SC-projection neurons^37,38^ (**Figure 7b**). This facilitates a better understanding of intrinsic SC circuitry and how convergent cortico-tectal projections onto zone-specific SC neurons influence the thalamic outputs^39,40^. Here, reconstructed neurons are superimposed above their corresponding SC zones which also receive inputs from visual-related cortical areas (VIS, RSP, PTLp). The LD-projecting SC. neurons had pyramidal-shaped cell bodies with dendrites that extended dorsally to the superficial optic layers and within the intermediate layers. Of the RE-projecting SC.m/cm neurons, some had dendrites that extended into superficial and intermediate layers, with others that extended into intermediate and deep layers. These neurons exhibited heterogeneous morphological properties consistent with the rat cell types that electrophysiologically respond to hyperpolarizing current pulses^41^. A subset of neurons extend dendrites horizontally, suggesting they integrate information within the same layer across zones. Other neurons extend vertically, suggesting they receive multimodal information integrated within multiple SC layers. Reconstructions of PF/MPT-projecting neurons in SC.cl/l zones overlap with inputs from somatic sensorimotor subnetworks. PF/MPT-projecting neurons in the SC.cl/l-ig/dg had wider dendritic arborizations than those in SC.m/cm.

We calculated a comprehensive battery of standard measurements including, number of bifurcations, contraction, width, and branch path length, to compare morphological features (**Figure 7c-d, Supplementary Figure 7**). See **Supplementary Table 7** for complete parameter list with definitions. Using all of the measured morphological parameters, principal component analysis (PCA) was run to reduce the dimensionality and create a 2D scatterplot (**Figure 7c**). The PCA shows the segregation of SC.m-PF/MPT neurons from all groups, including SC.m projecting neurons to both RE and LD. Neurons from SC.cm/cl/l projecting to PF/MPT are completely segregated from all SC.m projecting groups.

Statistically significant differences in several morphological features were detected across all pairwise comparisons (**Supplementary Table 6**). The parameters with greater loading values influencing the PCA are presented with whisker plots (**Figure 7d; Supplementary Figure 7**). Persistence-based neuronal feature vectorization framework was also applied to summarize pairwise differences between the nine groups of SC zone-specific projection neurons. The strength of the differences is presented as a gradient with white showing no difference and black showing the greatest differences (**Figure 7e**).

## DISCUSSION

The SC is a complex laminar structure implicated in spatial attention, orienting, navigation, defense, and decision-making^1,12,13,42,43^. As a major site of multisensory integration from visual, auditory, and somatosensory modalities, it is critical for animals to filter overwhelming sensory input, align brain-wide inputs, and guide behavior. Despite the wealth of literature on functional mapping studies in cats and monkeys, including newer studies in rodents, there remains a necessity for a comprehensive anatomical map of the mouse SC. Such a map would provide a foundational connectivity reference upon which functional hypothesis targeting discrete SC zones and cell types can be explored. During this resurgence of studies in the rodent SC, several studies provide entries into genetic and cell-type specific investigation of functional SC networks^36,44,45^. While individual cortico-tectal pathways are often separately studied, here, we comprehensively characterized all cortico-tectal projections based on the convergence and divergence of cortical fiber terminations across the rostrocaudal and mediolateral SC.

To construct the mouse cortico-tectal projectome, we first mapped all cortical inputs to the SC and identified four zone-specific delineations in the SC using a combined connectivity and computational neuroanatomic approach. In previous studies, we revealed the subnetwork organization of intracortical connectivity across the entire mouse neocortex^23^, and subsequently demonstrated how subnetwork projections are topographically conserved subcortically in the caudoputamen as the cortico-striatal projectome^24^. Thus, we hypothesized that cortico-tectal projections would conserve this subnetwork organization, and our results demonstrate that it is indeed conserved. Our analysis of cortico-tectal connectivity revealed new insights into the top-down cortical convergence of higher-order associative and prefrontal input with multimodal maps in the SC (**Figure 4**). The representation of sensory information shifts from medial SC.m and SC.cm zones which predominantly comprise visual and auditory associative subnetworks, centrolaterally into the SC.cl zone, and further lateral to the somatic sensorimotor subnetworks in SC.l. Furthermore, in addition to using cortico-tectal projections to delineate regional zones in SC, we analyzed the inputs and outputs of SC from thalamic, hypothalamic, midbrain and hindbrain. Medial associative subnetworks relaying visual, attention and spatial navigation information have integrated connectivity projections across all layers of SC.m and SC.cm (**Figure 8**).

**Figure 8.**
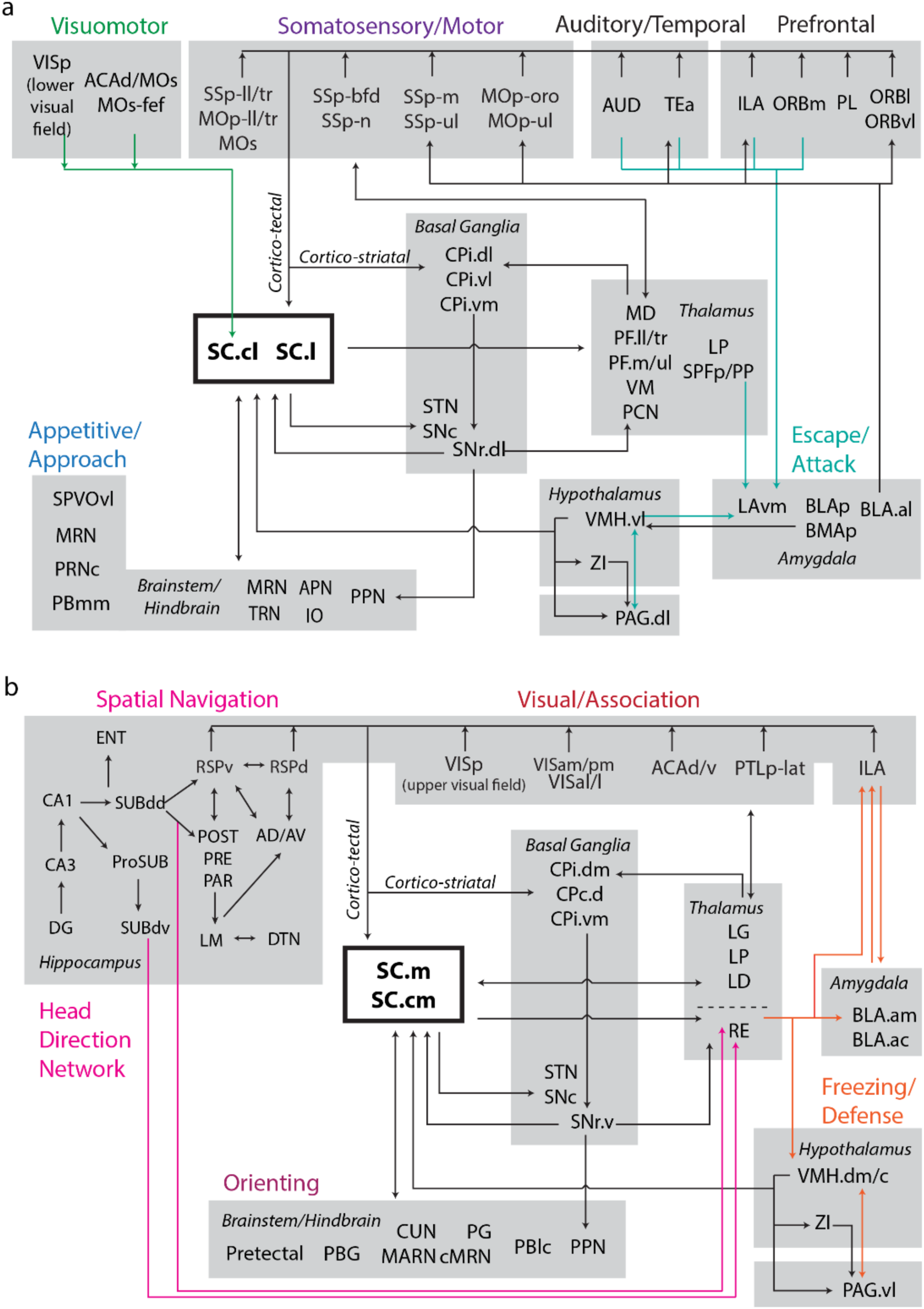
Subnetwork organization of SC zones. **a)** SC.m and SC.cm integration of visual information with spatial-related and head-direction hippocampal networks, attention/orienting regions, and freezing/defense regions. **b)** Integration of SC.cl and SC.l zones in visuomotor, somatic sensorimotor, escape and approach subnetworks (see text for detailed discussion). See **Supplementary Table 1** for complete abbreviation list.

Anatomical and functional studies show the SC interacts with the basal ganglia through connections with the thalamus and substantia nigra^29,46–48^. Our findings demonstrate topographically distinct and zone-specific ascending outputs to thalamic nuclei while the descending projections to the brainstem are also organized in alignment with SC zones. Each SC zone receives cortical information both directly through the cortico-tectal-thalamic projection pathway and indirectly through the multi-synaptic cortico-basal ganglia projection pathway. We selectively targeted RE, LD, and PF/MPT thalamic projection neurons from each SC zone and revealed significant distinctions in their morphometric features^49,50^. Cells projecting from SC.m superficial layers were more diverse than other zones, whereas cells from intermediate and deeper layers in SC.cm/cl had more similar features. Importantly, cells for this analysis were selected as representative thalamus projecting SC neurons and are not representative of the entire population of neurons within the four zones or layers. It is possible that brainstem projecting SC neurons display morphological features that differ from ascending thalamus projecting SC neurons. Nevertheless, our results suggest that functional analyses of SC neurons involving motor, sensory, and cognitive behaviors can also be addressed in the context of inputs and outputs from distinct modular networks. More studies are needed to characterize how multimodal cortical and subcortical information is integrated at single neuron resolution.

The columnar SC zones are also congruent with intrinsic collicular connectivity that integrates information across superficial, intermediate and deep layers, a principle of organization comparable to cortical columns within the neocortex^25,27,39,51^. Future studies can explore if these cortico-tectal zones reflect functional columns, and our work provides an anatomical foundation for hypothesis-driven investigations of cell-type specific functional behaviors. In humans, the SC is hypothesized to be a major locus of interest for potential therapeutic targets in treating hyper-responsivity and distractibility in attention-deficit hyperactivity disorder^5–8^. Understanding the neural networks of attention-related visual behaviors in mouse models has clinical significance in examining aberrant connections of neurodevelopmental disorders^4^.

### Functional hypotheses based on brain-wide anatomical subnetworks

#### Cortico-tectal subnetworks in navigation and goal-oriented behaviors

Studies on the postnatal development of the mouse cortex and SC reveal that the cortex must first codify multisensory cues formed from specific environmental contexts before they can effectively have a cortico-tectal influence on multisensory SC neurons^52^. This suggests that in addition to the array of innate responses to unconditioned sensory stimuli that the SC also has the capacity to integrate inputs from learned experiences to navigate and mediate goal-oriented responses via the basal ganglia. As described, RSP, PTLp-lat, and VIS prominently align in medial zones SC.m and SC.cm (and partially in SC.cl) that process distinct representations of visual information from the upper peripheral and central regions of the visual field^36,53^. RSP→SC presumably contributes to mediating spatial orientation and navigation, particularly through head-direction cells in RSPv integrating with the CA1dr/subiculum hippocampal network^45,54,55^. SC.m/cm in turn project to RE in thalamus which also contains head direction-sensitive cells^56^.

As environments vary during navigation, the overlapping inputs from ACA→SC may ontribute to facilitating attention during coordinated exploratory and evaluative response strategies. The ACA is implicated in guiding response selection and responding to the value in context, richness, and memory of an environment during exploratory and foraging behavior^57^. Our findings of the visuomotor subnetwork in SC.cl further support this zone as an analogous point of convergence for frontal eye field orienting behaviors in primates and cats^2^. As such, the widespread projections from ACAd/v to SC.cm, SC.cl and SC.l provide information regarding the contextual value of the environment that integrate with multisensory SC neurons processing input from virtually the entire visual, auditory, and somatotopic fields. Cells in the SC.cl-ig also aligned within acetylcholinesterase-rich SC compartments that project strongly to the MD thalamic nucleus which is also involved in frontal eye field processing^58–60^. By incorporating value-defined inputs across the SC, we can better understand the role of distinct SC zones in commanding eye movements and goal-oriented motor behaviors^61^.

#### Cortico-tectal subnetworks in approach/appetitive behaviors

The cortico-tectal lateral zones have functional implications in somatic sensorimotor movements that facilitate appetitive/approach behaviors^17,62^. SC.cl/l neurons receive inputs from MO/SS (orofacial, barrel field, upper limb) cortices, motor-related domains from thalamic nuclei (PF.mouth, PF.upper limb, VM, PCN), sexual approach behaviors (VMH.vl) and orofacial/upper limb brainstem and cerebellar inputs (**Figure 8**). These inputs provide evidence for SC.cl/l zones as distinct somatic sensorimotor subnetworks that mediate a subset of behaviors distinct from the more visually integrated range of behaviors mediated through SC.m/cm.

Our network analyses reveal the segregation of ventromedial prefrontal cortex (vmPFC) outputs from ORB and ILA regions to functionally specific SC.cl/l zones. By contrast to ACA in searching/foraging tasks, the vmPFC does not show any significant increase in activity during searching, but rather it responds to the values encoded between options chosen during searching^57^. This suggests that that vmPFC→SC provides prefrontal top-down input to command emotional and decision-making related responses. The sparse projections arising from ORBm/vl→SC.cl overlap with ACAd/MOs-fef→SC.cl inputs that target the lower nasal and temporal regions of the visual field represented in SC.cl, which is medially adjacent to the somatic sensorimotor subnetworks to facilitate spatial orientation near the ground. We observed that at least a subset of MOs/ACAd cortical projecting neurons (presumably pyramidal tract projection neurons) generate collateral projections to both the SC.cl and CPc.d (**Supplementary Figure 6c**). As such, each zone of the SC receives cortical information both directly through the cortico-tectal projection pathway and indirectly through the multi-synaptic cortico-striato-nigra-tectal projection pathway.

The lateral PFC projections from ORBl→SC.l inputs align with barrel-field and oropharyngeal subnetworks in SC.l, thereby facilitating decision-making about relevant stimuli proximal to the ground^62,63^. The combination of these limbic system inputs may facilitate the integration of valence and motivated emotional responses in social and appetitive behaviors^64^.

#### Cortico-tectal subnetworks in defensive/aggressive behaviors

The cortico-tecto-thalamic subnetwork has intimate connectivity with hypothalamic and amygdalar structures to coordinate activation of fear-related circuitry^60,64,65^. Within the medial cortico-cortical subnetworks, ORB is interconnected with ACAv, ACAd/MOs, and PTLp-med (internal body map), which project to the SC.cl/l zones, suggesting a coordination of motor outputs for the whole body to prompt escape and attack reactions (**Figure 8**). Notably, the SC.m and SC.cm deep layers also receive dense input from the dorsomedial part of the VMH (VMH.dm/c) known to evoke innate defensive responses and aggressive responses to predators^66–68^. These medial SC zones send outputs to the RE, CM, and IAM shown to be activated during arousal and the presence of visual threats^69,70^. Our results also reveal an interesting divergent ILA→SC.m/l projection pathway that is consistent with the role of ILA in mediating fear extinction, risk assessment and evaluation to threat situations, particularly to looming stimuli and predators^17,18,57,71^.

## CONCLUSION

Overall, these results refine previous studies observing sensory topographic distribution patterns within the SC, and provide a comprehensive understanding of how all higher-order cortical inputs are integrated. To the best of our knowledge, this map of the cortico-tectal projectome is the first in the mouse model and will thus serve as a useful cross-species reference^15,16,72^. We have shown the mouse SC can be subdivided into four columnar zones with distinct connectivity that correlate with functional subnetworks. For each zone, we characterized the input/output organization and provide several working hypotheses to explore their functional implications. This study advances our understanding of SC organization within brain-wide subnetworks, and is a useful resource for investigating the role of cortico-tectal networks in disease models. Ultimately, the cortico-tectal projectome provides an anatomical framework for understanding how individual SC cell-types integrate convergent input from multiple brain areas and functional modalities for the control of attention and other goal-directed behaviors.

## METHODS

### Mouse Connectome Project methodology

Anatomical tracer data was generated as part of the Mouse Connectome Project (MCP) within the Center for Integrative Connectomics (CIC) at the University of Southern California (USC) Mark and Mary Stevens Neuroimaging and Informatics Institute. MCP experimental methods and online publication procedures have been described previously^23,24,54,73^. We systematically and carefully mapped neuronal connectivity of every cortical structure to determine their input connectivity to the superior colliculus (SC) (for complete injection site list, see **Supplementary Table 2**). We used multiple fluorescent tracing strategies with a combination of classic tract-tracing and viral tracing methods. First, we used a triple anterograde tracing approach with individual injections of PHAL and GFP- and tdTomato-expressing adeno-associated viruses. Additionally, to investigate the convergence or divergence of axonal fiber pathways either into or out of the SC, we used a double co-injection approach that injects two different tracer cocktails each containing one anterograde and one retrograde tracer to simultaneously visualize two sets of input/output connectivity. Data generated is published online as part of the Mouse Connectome Project (www.MouseConnectome.org).

#### Animal subjects

All MCP tract-tracing experiments were performed using 8-week old male C57BL/6J mice (n=60; Jackson Laboratories). Mice had ad libitum access to food and water and were pair-housed within a temperature- (21-22°C), humidity- (51%), and light- (12hr: 12hr light/dark cycle) controlled room within the Zilkha Neurogenetic Institute vivarium. All experiments were performed according to the regulatory standards set by the National Institutes of Health Guide for the Care and Use of Laboratory Animals and by the institutional guidelines described by the USC Institutional Animal Care and Use Committee.

#### Tracer injection experiments

The MCP uses a variety of combinations of anterograde and retrograde tracers to simultaneously visualize multiple anatomical pathways within the same Nissl-stained mouse brain. Triple anterograde tracing experiments involved three separate injections of 2.5% *phaseolus vulgaris* leucoagglutinin (PHAL; Vector Laboratories, Cat# L-1110, RRID:AB_2336656), and adeno-associated viruses encoding enhanced green fluorescent protein (AAV-GFP; AAV2/1.hSynapsin.EGFP.WPRE.bGH; Penn Vector Core) and tdTomato (AAV1.CAG.tdtomato.WPRE.SV40; Penn Vector Core). An anterograde tracer of 5% biotinylated dextran amine (BDA; Invitrogen) was used in some cases. Retrograde tracers included cholera toxin subunit B conjugates 647, 555 and 488 (CTb; AlexaFluor conjugates, 0.25%; Invitrogen), Fluorogold (FG; 1%; Fluorochrome, LLC), and AAVretro-EF1a-Cre (AAV-retro-Cre; Viral Vector Core; Salk Institute for Biological Studies). To provide further details on specific connectivity patterns, we also performed quadruple retrograde tracer, and rabies/PHAL experiments. Quadruple retrograde tracer experiments involved four different injections sites receiving a unique injection of either 0.25% CTb-647, CTb-555 CTb-488, 1% FG, or AAV-retro-Cre.

To trace the connectivity of projection defined neurons, a Cre-dependent anterograde tracing strategy was employed, which involved the delivery of Cre to projection neurons of interest via an injection of AAVretro-EF1a-Cre into their target regions. Next, AAV1.CAG.FLEX.eGFP.WPRE.bGH and AAV1.CAG.FLEX.TdTomato.WPRE.bGH (Addgene; plasmid ID #s 51502, 51503) were delivered to the different Cre-expressing SC neuronal populations. Finally, to retrieve the morphological information from different tectal projection neurons, Gdel-RV-4tdTomato^49^, and Gdel-RV-4eGFP^50^ injections were made into two different downstream targets of the neurons. All cases used in this study are listed in **Supplementary Table 2**. No statistical methods were used to pre-determine sample sizes, but our sample sizes are similar to those reported in previous publications. In most cases, anterograde tracing results are cross validated by retrograde labeling injections at anterograde fiber terminal fields and vice versa.

#### Stereotaxic surgeries

On the day of experiment, mice were deeply anesthetized and mounted into a Kopf stereotaxic apparatus where they are maintained under isofluorane gas anesthesia (Datex-Ohmeda vaporizer). For triple anterograde injection experiments, PHAL was iontophoretically delivered via glass micropipettes (inner tip diameter 24-32μm) using alternating 7sec on/off pulsed positive electrical current (Stoelting Co. current source) for 10min, and AAVs were delivered via the same method for 2 min (inner tip diameter 8-12μm). For anterograde/retrograde coinjection experiments, tracer cocktails were iontophoretically delivered via glass micropipettes (inner tip diameter 28-32μm) using alternating 7sec on/off pulsed positive electrical current (Stoelting Co. current source) for 5 (BDA or AAV/FG) or 10 min (PHAL/CTB-647). For quadruple retrograde tracing experiments, 50nl of retrograde tracers were individually pressure-injected via glass micropipettes at a rate of 10nl/min (Drummond Nanoject III). Rabies injections of 10nL were individually pressure-injected via glass micropipettes (inner tip diameter 8-12μm). All injections were placed in the right hemisphere. Injection site coordinates for each surgery case are in **Supplementary Table 2** based on the (ML, AP, DV) coordinate system from the 2008 Allen Brain Reference coronal atlas^19^. Following injections, incisions were sutured, and mice received analgesic pain reliever and were returned to their home cages for recovery.

#### Histology and immunohistochemical processing

After 1-3 weeks post-surgery, each mouse was deeply anesthetized with an overdose of Euthasol and trans-cardially perfused with 50ml of 0.9% saline solution followed by 50ml of 4% paraformaldehyde (PFA, pH 9.5). Following extraction, brain tissue was post-fixed in 4% PFA for 24-48hr at 4°C. Fixed brains were embedded in 3% Type I-B agarose (Sigma-Aldrich) and sliced into four series of 50μm thick coronal sections using a Compresstome (VF-700, Precisionary Instruments, Greenville, NC) and stored in cryopreservant at -20°C. For double coinjection experiments, one series of tissue sections was processed for immunofluorescent tracer localization. For PHAL or AAVretro-EF1a-Cre immunostaining, sections were placed in a blocking solution containing normal donkey serum (Vector Laboratories) and Triton X (VWR) for 1 hr. After rinsing in buffer, sections were incubated in PHAL primary antiserum (1:100 rabbit anti-PHAL antibody (Vector Laboratories Cat# AS-2300, RRID:AB_2313686)) or AAVretro-EF1a-Cre primary antiserum (1:100 mouse-anti-Cre Recombinase antibody, clone 2D8, Millipore Sigma Cat# MAB3120) mixed with donkey serum, Triton X, in KPBS buffer solution for 48-72 hours at 4°C. Sections were then rinsed again in buffer solution and then immersed in secondary antibody solution (donkey serum, Triton X, and 1:500 donkey anti-mouse IgG conjugated with Alexa Fluor 647 (Life Technologies Cat# A-31571), or 1:500 donkey anti-rabbit conjugated with 647 (Jackson ImmunoResearch Labs Cat# 715-165-151, RRID:AB_2315777) for 3 hrs. BDA immunofluorescence was visualized using a 647- or 568-conjugated Streptavidin. Finally, all sections were stained with Neurotrace 435/455 (Thermo Fisher Cat# N21479) for 2-3 hours to visualize cytoarchitecture. After processing, sections were mounted onto microscope slides and cover slipped using 65% glycerol.

### Imaging and post-acquisition processing

Complete tissue sections were scanned using a 10X objective lens on an Olympus VS120 slide scanning microscope. Each tracer was visualized using appropriately matched fluorescent filters and whole tissue section images were stitched from tiled scanning into VSI image files. For online publication, raw images are corrected for correct left-right orientation and matched to the nearest Allen Reference Atlas level (ARA)^19^. An informatics workflow was specifically designed to reliably warp, reconstruct, annotate and analyze the labeled pathways in a high-throughput fashion through our in-house image processing software Connection Lens (**Figure 1d**). Tissue sections from each analyzed case were assigned and registered to a standard set of 10 corresponding ARA levels ranging from 84-102 (all images shown in this manuscript are from unwarped, unregistered VSI images). Threshold parameters were individually adjusted for each case and tracer. Adobe Photoshop was used to correct conspicuous artifacts in the threshold output files. Each color channel was brightness/contrast adjusted to maximize labeling visibility (Neurotrace 435/455 is converted to brightfield) and TIFF images were then converted to JPEG file format for online publication in the Mouse Connectome Project iConnectome viewer (www.MouseConnectome.org).

#### Assessment of injection sites and fibers of passage

All cortical injection cases included in this work are, in our judgment, prototypical representatives of each cortical area. We have previously demonstrated our targeting accuracy with respect to injection placement, our attention to injection location, and the fidelity of labeling patterns derived from injections to the same cortical location. In the current report, we also demonstrate our injection placement accuracy and the consistent labeling resulting from injections placed in the same cortical areas (see Supplementary Methods in Hintiryan et al., 2016 for details). Anterograde cortico-tectal labeling patterns were cross-validated with retrograde injections in the SC to backlabel distribution of cortical inputs (Supplementary Figs. 2b, 3c). A combination of labeling produced from both anterograde and retrograde cases across cortex and SC was used to generate the input/output connectome diagram (**Figure 5**).

An important concern when employing automated analysis of connectivity is the issue of fibers of passage getting annotated as functional connections when in fact no synapses exist. Pathways devoid of synapses can produce bright labeling that gets annotated as positive pixels. This is especially relevant with regard to cortico-tectal fibers that travel from the rostral direction through the SC onto a caudal SC termination site. For example, ARA 86 level exhibited more labeling from fibers of passage entering rostrally from both sensorimotor and higher-order cortical areas such as MOp/MOs (**Figure 2a**), PTLp-med (**Figure 3a**) and RSPv-rostral (**Supplementary Figure 3a**). Some fibers appear to have evident terminal boutons, while others do not. So as to not introduce subjective confounds across the data, those pixel values remained as part of the computational analysis.

#### Defining the SC custom atlas and delineating SC zones

The workflow was applied toward analyzing projections from 86 representative cortical injection cases to sections with characteristic labeling across the far rostral SC (ARA 86), rostral SC (ARA 90), middle SC (ARA 96) and caudal SC (ARA 90). Atlas level 86 is the furthest rostral where notable fiber terminations were observed. The labeling at atlas level 90 (−3.68 mm from bregma) was the target of rich projections from all cortical areas, representative of all cortico-tectal projections, and exhibited the most segregation of labeling in terms of unique zones. Therefore, it was selected as the representative section for the rostral SC. The SC at level 86 (−3.28mm from bregma) was the furthest rostral level we could use for analysis that expressed prominent terminals from a subset of cases. Therefore, it was selected as the representative section for the far rostral SC. The SC at atlas levels 96 (−4.28mm from bregma) and 100 (−4.65mm from bregma) displayed distinguishable labeling with variable degrees of layer- and zone-specific cortical inputs and far fewer discrete termination fields compared to the rostral SC and were therefore selected to represent the middle and caudal SC, respectively.

We overlapped reconstructed cortico-tectal fibers from the 86 cases to each custom SC level (86, 90, 96, and 100), and determined the boundaries of the four zone delineations based on the average patterns generated. Some cortico-tectal patterns naturally distributed across more than one zone, though the general boundaries were still consistent within the four zones. As representative examples, anterograde AAV injections into the right hemispheres of PTLp (red), VISam (orange), ACAd (green) and MOp (purple) terminate with different laminar and regional patterns in SC (ARA 90) (**Figure 1b**). PTLp produced ipsilateral terminals in the intermediate layers of SC, delineating the medial zone (SC.m). VISam produced tiered terminals in the SC superficial and intermediates layers adjacent to SC.m, delineating the centromedial zone (SC.cm). ACAd produced tiered terminals in the SC intermediate and deeper layers, laterally adjacent to the VISam terminal field, delineating the centrolateral zone (SC.cl). MOp produced terminals targeting the far lateral SC, delineating lateral zone (SC.l). Together, these representative cortico-tectal projections reveal layer-specific terminals distributed across four distinct zones along the medial-lateral axis of SC.

#### Color coding, data visualization, and proportional stacked bar charts

The same color scheme within SC zones was used for consistency of reference throughout the diagrams: SC.m (red), SC.cm (orange), SC.cl (green), SC.l (purple). In the topographic map of the brain dorsal view (**Figures 2b, 3c, 6e**), cortical areas were color-coded based on their SC zone target.

Stacked bar chart for visualization of proportion of labeling across each SC zone (*x* axis) from each cortical ROI (*y* axis). Values represent proportion of pixel density for the selected ROI (n=1 per cortical area) distributed across each SC zone across all layers. The chart facilitates an overview representation of how cortico-tectal projections preferentially target specific zones and how they compare across cortical groups. For complete breakdown of values for zone- and layer-specific proportion of label bar charts, see **Supplementary Tables 3-4**.

#### Community detection and connectivity matrices

We further analyzed the annotation data to objectively identify groups of cortical injection sites that send converging input within different SC zones. To perform this final stage of the neuroinformatics pipeline, we first built an adjacency matrix out of our annotation data. The graph structure of the data is relatively simple: nodes and connections are organized as a multi-tree with two levels: the cortex and the four SC zones. We performed community detection (modularity maximization) on ROI annotated data. The overlap annotation per each case within a group was aggregated into a single matrix. Overlap refers to a convergence of terminal fields within the SC between cortical source areas. The overlap value between a source and target ROI is the ratio of common labeling among the two ROIs to total source labeling^24^. Once the aggregated matrix was constructed, we further normalized so that the total labelling across each injection site (typically close in the first place) was adjusted to equal with the injection site featuring maximum total labeling. On this normalized matrix we applied the Louvain community detection algorithm at a single scale (gamma 1.0) to the data and identify clusters of injection sites with similar SC termination fields.

As the result of this greedy algorithm is non-deterministic, we performed 100 separate executions, and subsequently calculated a consensus community structure to characterize the 100 executions as a single result. Given the modularity of the data (that is, highly topographic labeling), a modularity optimization algorithm like the Louvain was well suited. However, the element of randomness makes it probable that the algorithm will reveal a different community structure over multiple runs. To mitigate this issue, the algorithm was run 1,000 times. The community structure that emerged most often, which we defined as the community structure mode (borrowing from statistics) is reported. A mean and s.d. for the number of communities that was detected across the 1,000 runs was also computed. Subsequently, to aid visualization we employed the community structure to re-order and color code an adjacency matrix such that connections were placed close to the diagonal. An accompanying color-coded SC illustrates the spatial arrangement of the communities/zones. The code for generating the illustrative color-coded SC employed a ‘winner takes all’ reconstruction of community terminal fields. For each cell, the method compared overlap data from across injection sites and colored according to the community with the greatest quantity of labeling.

For matrix visualization, we applied an algorithm to modularize ROIs based on community assignment, as well as prioritize connections along the diagonal. Such a visualization provides a high-level overview of the connectivity (**Figure 4**). Community coloring provides more detail by visually encoding segmentation by community, and ultimately, injection site (anterograde visuals). To carry out this process, we developed software to programmatically march through each segmented pixel per each image. Using ROI name, the algorithm looked up the corresponding community assigned during the consensus community step. Using a table containing a color assigned (by the authors) to each injection site, the algorithm retrieved the injection site associated with the community and colored the pixel with the corresponding injection site color value. A subsequent step took advantage of the fact that each pixel was assigned to only a single community by aggregating all colorized images corresponding to a given atlas level into a single representative image.

### 3D Workflow

To assess whether thalamic-projection neurons in the four SC zones were morphologically, representative neurons in each zone were labeled via a G-deleted-rabies injection in the LD (for SC.m/cm), in the RE (for SC.m/cm), and dorsal PF (**Figure 7a**). One week was allowed for tracer transport following injections, after which the animals were perfused. The SHIELD clearing protocol was used for the 3D tissue processing workflow^74,75^, followed by neuronal reconstructions and statistical analyses^71^.

#### 3D tissue processing

Mice were transcardially perfused with cold saline and SHIELD perfusion solution. The brains were extracted and incubated in the SHIELD perfusion solution at 4°C for 48 hours. The SHIELD perfusion solution was replaced with the SHIELD OFF solution and tissues were incubated at 4°C for 24 hours. The SHIELD OFF solution was replaced with the SHIELD ON solution and the tissues were incubated at 37°C for 24 hours. The whole brain was cut into 400µm sections and were cleared in the SDS buffer at 37°C for 72 hours. The sections were then washed three times with KPBS and incubated in KPBS at 4°C for 24 hours.

#### 3D imaging protocol

Sections were mounted and coverslipped onto 25×75×1mm glass slides with an index matching solution 100% (EasyIndex, LifeCanvas Technologies, #EI-Z1001). Sections were imaged with a high-speed spinning disk confocal microscope (Andor Dragonfly 202 Imaging System, Andor an Oxford Instruments Company, CR-DFLY-202-2540). 10x magnification (NA 0.40, Olympus, UPLXAPO10X) was used to acquire an overview after which 30x magnification (NA 1.05, Olympus, UPLSAPO30xSIR) was used to image through the SC ipsilateral to the injection site at 1 µm z steps. Sections were imaged with four excitation wavelengths (nm): 405 (blue Nissl background), 488 (green for Rabies), 561 (red for Rabies) with respective emission detection wavelengths of 450, 525, and 600.

#### 3D reconstructions, visualizations, and analysis of neuronal morphology

Manual reconstruction of the neurons was performed using Aivia (version.8.8.1, DRVision) (**Figure 7**), and geometric processing of neuron models, including smoothing, was performed using the Quantitative Imaging Toolkit (QIT) NeuronTransform and NeuronFilter modules^76^ (also available at http://cabeen.io/qitwiki). neuTube was used to facilitate visualizations of reconstructions^77^. All reconstructions are being made freely available in the Dong archive of www.NeuroMorpho.Org^22^. Although we wanted to compare projection neurons from each SC zone to each thalamic area, neurons from each injection were spread across two to four zones. This allowed us to compare SC neurons in neighboring zones that project to the same thalamic nucleus, as well as those that project to a different thalamic nucleus. Three groups were assigned from the RE rabies injected case based on location of retrogradely labeled cells in SC: RE-SC.m (n=23), RE-SC.cm (n=27), and RE-SC.cl (n=4). Two groups were assigned form the LD case: LD-SC.m (n=4), and LD-SC.cm (n=4). Four groups were assigned from the PF/MPT case: PF/MPT-SC.m (n=10), PF/MPT-SC.cm (n=6), PF/MPT-SC.cl (n=9), PF/MPT-SC.l (n=5).

#### Statistical analyses on groups of reconstructions

To obtain an overall view of the dendritic morphology of SC projection neurons located in the SC.m, SC.cm, and SC.cl, we applied the classic Sholl analysis using Fiji ImageJ. Quantitative morphological parameters characterizing arbor morphology were obtained from L-Measure^38^ and statistical analyses were performed using the R computing environment (RStudio Version 1.1.463). Standard measurements of dendritic morphology included height, width, height/width ratio, length, radius of neuronal shell (Euclidean distance), number of stems attached to the soma, number of bifurcations, number of branches, number of tips, fragmentation, contraction, local bifurcation amplitude angle, remote bifurcation amplitude angle, local bifurcation up/down angle, remote bifurcation up/down angle, fractal dimension, centrifugal branch order, partition asymmetry and mean synaptic distance (path distance). Using all of the measured morphological parameters, principal component analysis (PCA) was run to reduce the dimensionality and create a 2D scatterplot. The PCA shows the segregation of SC zone-specific neurons based on the measured features (**Figure 7c**).

Two-sided pairwise Wilcoxon rank sum tests were performed, and the parameters that survived FDR correction for multiple testing with a p-value <0.05 are reported. Complete list of output p-values can be found in **Supplementary Table 7**. Significant differences were detected across several pairwise parameters. Examples of the significant group differences are presented with whisker plots (**Figure 7e**; **Supplementary Figure 7**). Complete list of calculated whisker-plot values (minimum, 1^st^ quadrant, median, mean, 3^rd^ quadrant, and maximum) for each parameter can be found in **Supplementary Table 6**.

We characterized our data using an alternative tool that uses topological data analysis that retains potentially more morphological information. Persistence-based neuronal feature vectorization framework was also applied to summarize pairwise differences between the nine groups of SC zone-specific projection neurons. The strength of the differences is presented as a gradient with white showing no difference and black showing the greatest differences (**Figure 7e**). Our experiments used code provided by the authors online, where we first computed persistence diagrams using NeuronTools (https://github.com/Nevermore520/NeuronTools) and then computed inter-neuron distances using the Wasserstein metric (https://bitbucket.org/grey_narn/geom_matching/src). The results were visualized in matrix plots created using R (**Figure 7; Supplementary Figure 7**). Similar to the PCA analysis, the data showed that groups of projections neurons in SC.m, SC.cm, SC.cl, and SC.l all differed from one another, but the largest differences were between SC.m neurons (projecting to MPT from superficial layers) compared to those in each other zone (**Figure 7**).

### Code Availability

All custom code that we developed for data analysis is available at https://git.ini.usc.edu/lgarcia/SC. Code used for other data analyses are freely accessible. For the Louvain algorithm implementation, the Brain Connectivity Toolbox (BCT) was employed, available at https://sites.google.com/site/bctnet/ for matrix visualization. The grid_communities algorithm for matrix visualizations are available at https://github.com/aestrivex/bctpy. Geometric processing of neuron models was performed using the Quantitative Imaging Toolkit (QIT), available here http://cabeen.io/qitwiki. We also computed persistence diagrams using NeuronTools (https://github.com/Nevermore520/NeuronTools) and then computed inter-neuron distances using the Wasserstein metric (https://bitbucket.org/grey_narn/geom_matching/src. In addition, we have used several open-source Python packages: scipy (v0.17.0), matplotlib (v1.5.1), ipython (v2.4.1), ipython-genutils (v0.2.0), jupyter (v1.0.0), jupyter-client (v5.2.3), jupyter-console (v5.2.0), jupyter-core (v4.4.0), pandas (v0.23.3), sympy (v1.2), nose (v1.3.7), pydot (v1.2.3), bctpy (v0.5.0), pyflakes (v1.1.0), pandas (v0.23.3), cycler (v0.9.0), plotly (v3.0.0).

## DATA AVAILABILITY

Anatomical tracer image data is available through our iConnectome viewer as part of the Mouse Connectome Project at USC (http://www.mouseconnectome.org). Additional data that support the findings of this study that are not readily accessible online are available from the corresponding author upon reasonable request.

## ACKNOWLEDGMENTS

This work was supported by NIH/NEI F31 EY029569 (N.L.B.), NIH T32 MH111360 (N.L.B), NIH/NIMH R01 MH094360 (H.-W.D.), NIH/NIMH U01 MH114829 (H.-W.D.), NIH/NCI U01 CA198932-01 (H.-W.D), NIH/NIMH U19 (JE/EC) and NIH/NIMH U19 MH114821 (J.H./P.A.).

## AUTHOR CONTRIBUTIONS

N.L.B., M.S.B. and H.-W.D. conceived, designed and managed the project. N.L.B. and H.-W.D. wrote the manuscript. N.L.B. performed manual analysis of all raw image data, including the connectivity annotation, created the SC custom atlas, and prepared figures for publication. I.B. and L.G. wrote the code for computational network analysis. L.G. and N.L.B. designed matrices, stacked bar charts, and anterograde maps. N.L.B., N.K., and M.F. registered cases on Connection Lens for analysis. N.L.B., M.S.B. and H.-W.D. constructed the neural networks diagrams. N.L.B., L.G., L.G., M.B., M.Y.S., N.F.F., S.A, and C.C. performed stereotaxic surgeries and tissue processing to generate anatomical connectivity data. N.K., B.Z., and N.L.B. processed and designed tissue clearing and 3D imaging protocols. L.K. traced all 3D reconstructed neurons. L.K. and G.A.A. performed statistical analysis of morphological data. I.R.W. provided rabies viruses. S.Y. managed the iConnectome website and created online informatics and visualization tools. K.C. helped implement the SC custom atlas on Outspector. N.K., M.F., D.J., D.L., T.B., and S.U. performed image processing for image data uploaded to the iConnectome viewer. M.S.B., B.Z., G.A.A., H.H., and A.S. offered constructive guidance for the manuscript edits.

## Supplementary Materials

**Supplementary Figure 1.**
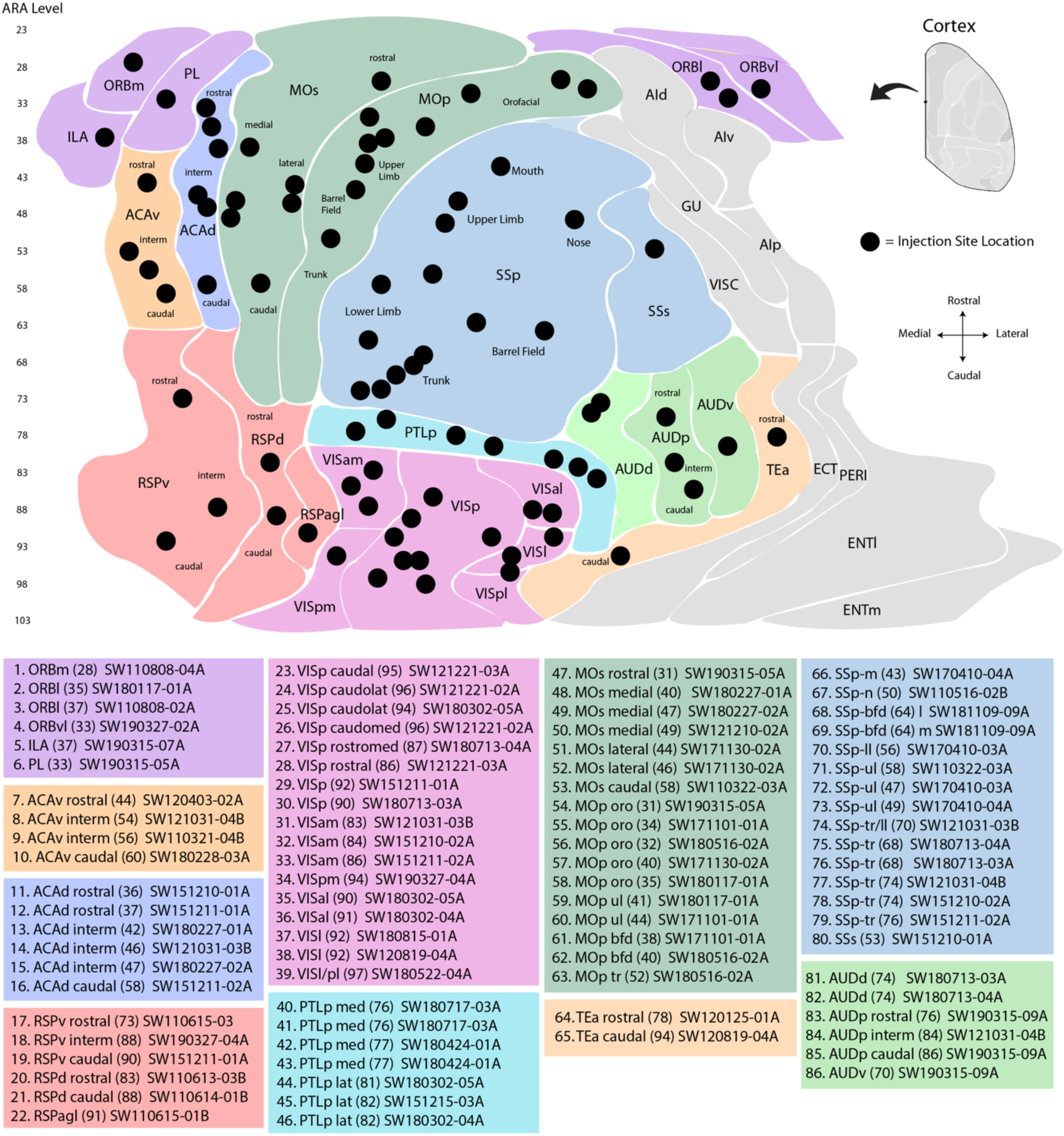
Flat map of mouse cortex with injection sites names. Injection site map was generated by expanding the length of each cortical area from the coordinate framework on the Allen Institute Adult Mouse Brain Atlas. Filled circles indicate location of injection site. List of injection sites, coordinates, and location can be cross-referenced by index number with **Supplementary Table 2**.

**Supplementary Figure 2.**
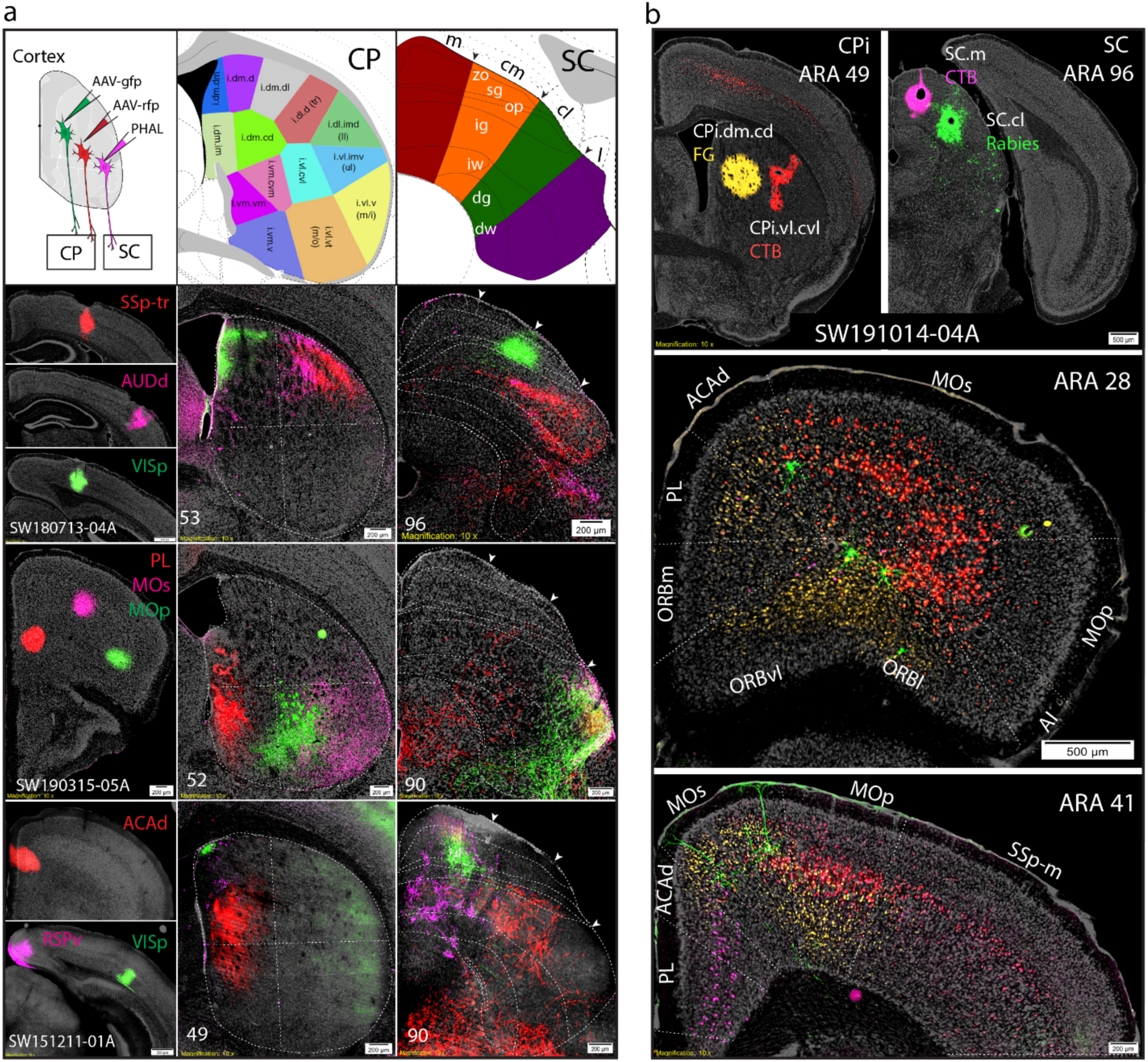
Conserved organization between sensory related cortico-striatal domains and cortico-tectal zones. **a)** Three separate triple anterograde cases into different cortical areas send topographic projections to corresponding caudate putamen (CP) domains and SC zones. **b)** Quadruple retrograde case into CPi.dm.cd (FG), CPi.cl.cvl (CTB-555), SC.m/cm (CTB-647) and SC.cl (Rabies-GFP). Distribution of retrogradely labeled cells across cortex.

**Supplementary Figure 3.**
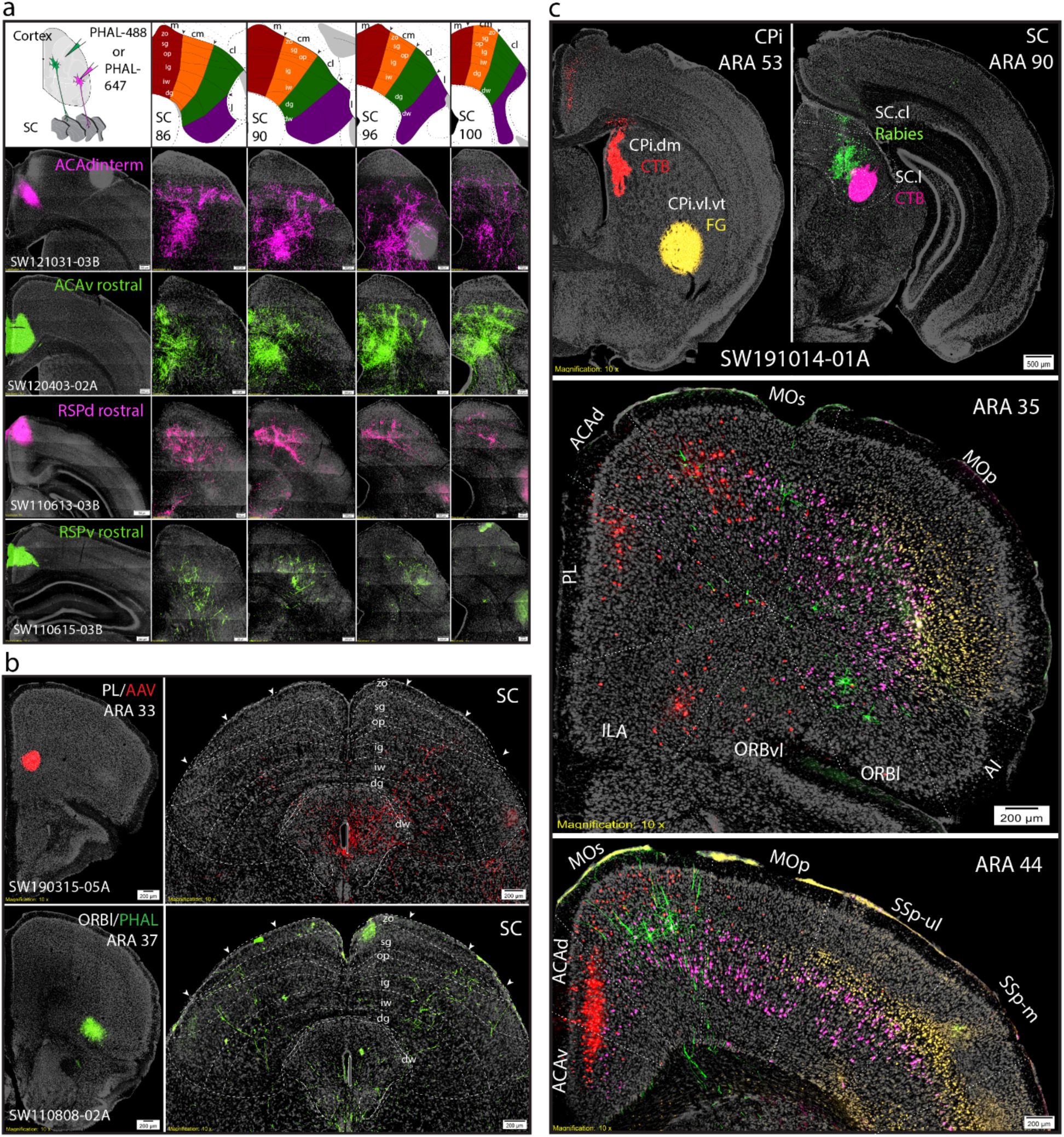
Higher-order distribution arrays. **a)** Data array for individual cases from higher order projections to SC. **b)** Two cases with single anterograde injections into PL (AAV) and ORBl (PHAL), and their respective bilateral projections predominately to SC.cl and SC.l zones. **c)** Quadruple retrograde case into CPi.dm (CTB), CPi.cl.vt (FG), SC.cl (Rabies-GFP), and SC.l (CTB). Distribution of retrogradely labeled cells across cortex.

**Supplementary Figure 4.**
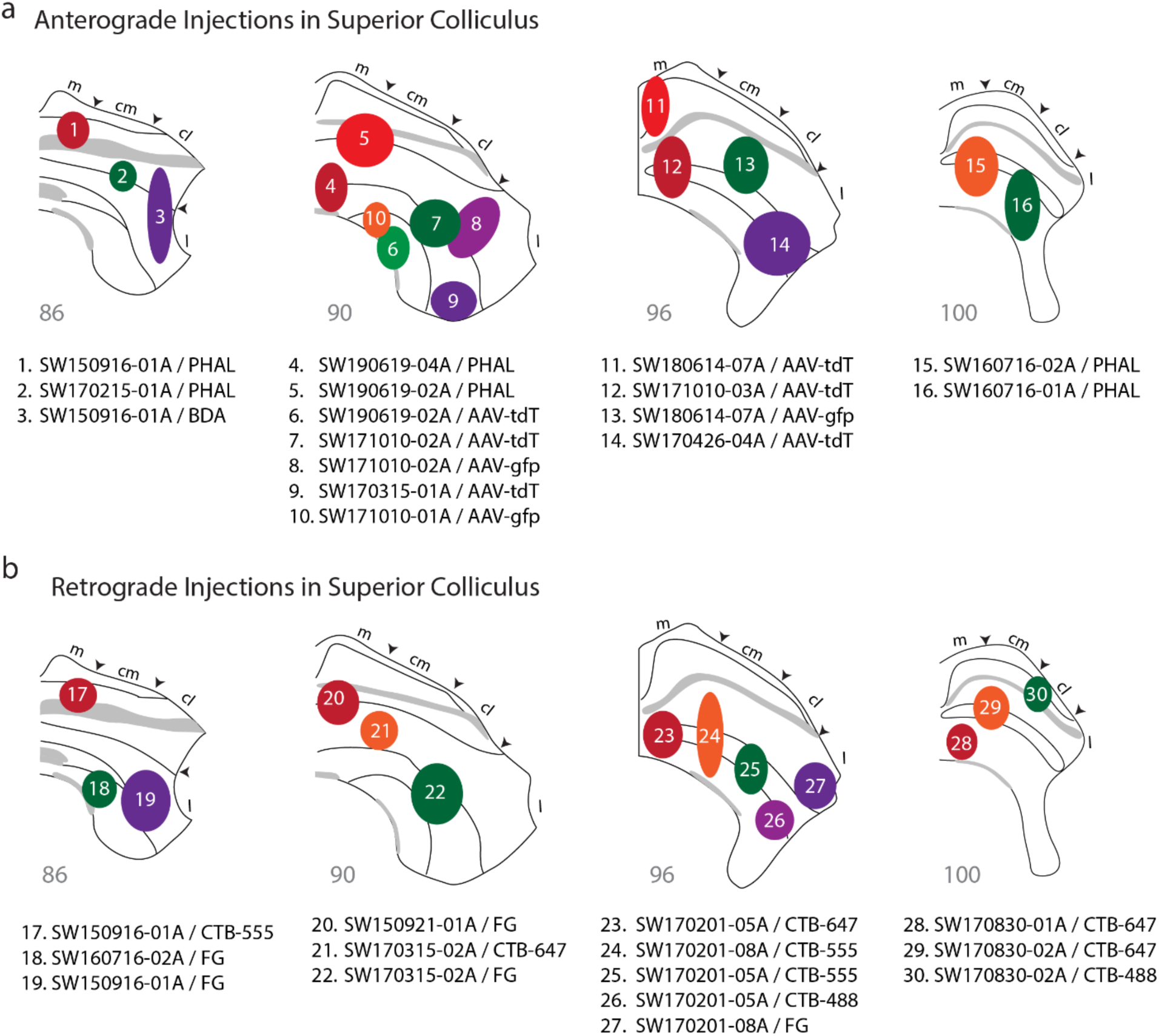
Superior colliculus injection site map. **a)** Anterograde injection sites in superior colliculus. **b)** Retrograde injection sites. List of injection sites, coordinates, and location can be cross-referenced by index number with **Supplementary Table 2**. *Abbreviation list: AAV, adeno-associated virus; AAV-tdT, AAV-tdTomato; AAV-gfp, AAV-green fluorescent protein; BDA, biotinylated dextran amine; CTB, cholera toxin subunit B conjugates (488, 555, 567); FG, Fluorogold; PHAL, phaseolus vulgaris leucoagglutinin.*

**Supplementary Figure 5.**
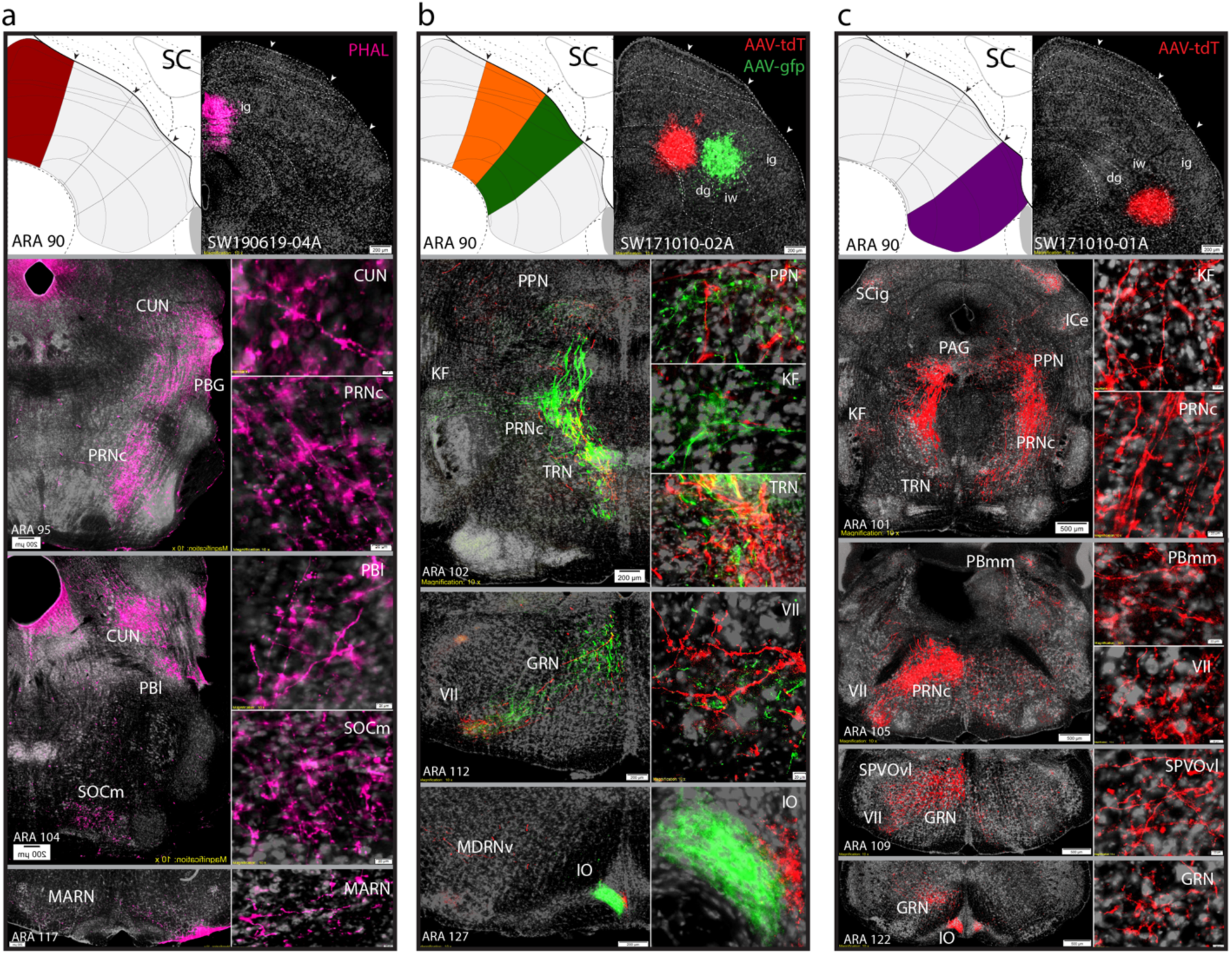
Downstream projections from distinct SC zones to brainstem regions. **a)** SC.m-ig ipsilateral projections. **b)** SC.cm-ig/iw and SC.cl-ig/iw injections target similar brainstem regions, with segregated outputs in IO. **c)** SC.l-dg/iw projections. An anterograde map of reconstructions distributed across the entire brain will be available upon publication.

**Supplementary Figure 6.**
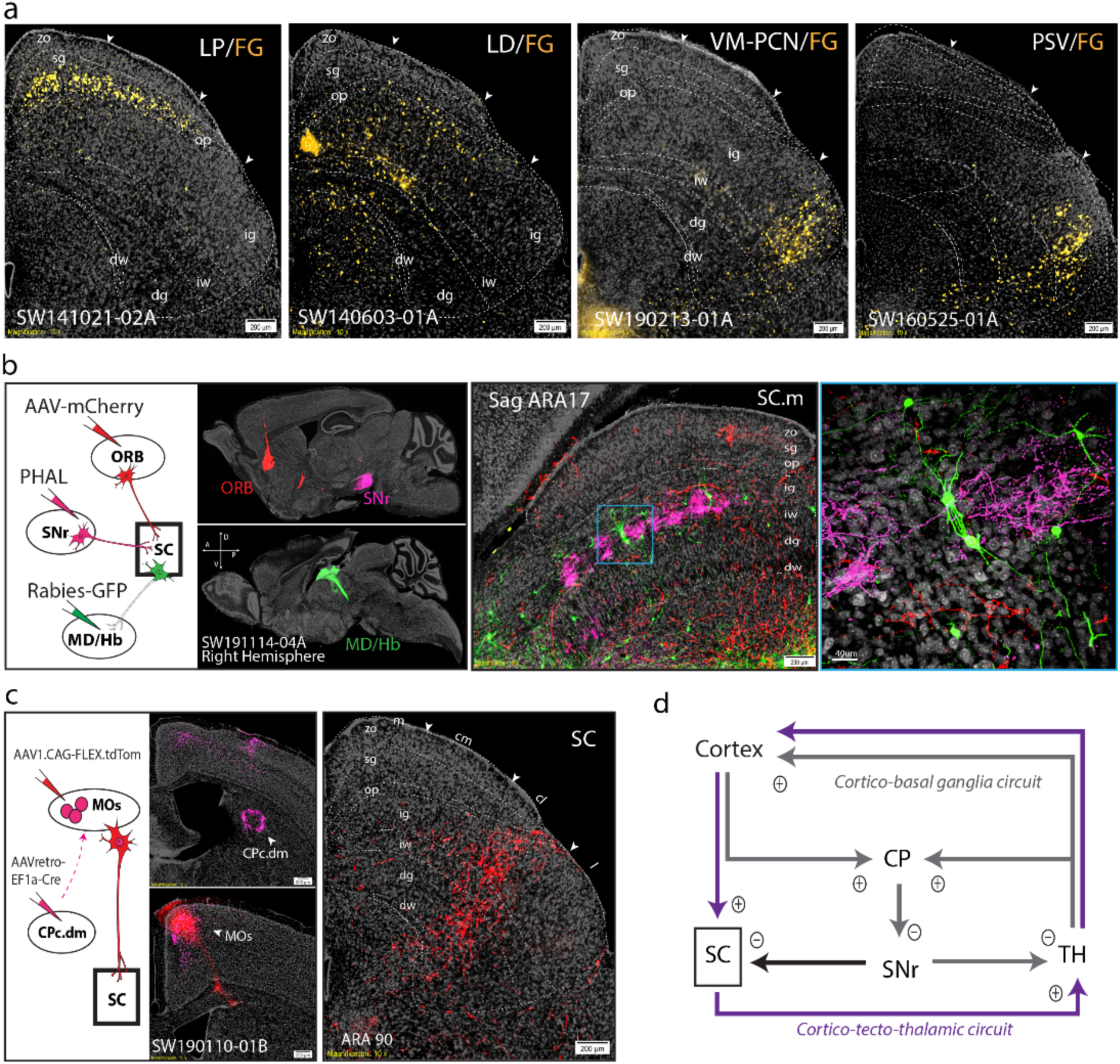
Brain-wide network connectivity. **a)** Three cases with retrograde labeling from FG injections in thalamic and brainstem nuclei. Left: LP injection labeled cells in sg layer predominantly across SC.m/cm. Left middle: LD injection labeled cells across ig/iw/dg layers predominantly in SC.m/cm. Right middle: VM-PCN injection labeled cells exclusively in SC.l. Right: PSV injection labeled cells in SC.l. **b)** Anterograde projections from ORB (AAV-mCherry) and from SNr (PHAL) synapse onto the same Rabies-GFP labeled SC neurons that project to MD/Hb in the thalamus. This circuit shows the cortico-tectal-thalamic loop that integrates with the SNr outputs of the basal ganglia. **c)** A combinatorial tract tracing method using AAVretro-Cre injection (in CPc.d) and Cre-dependent AAV-mCherry (in MOs/ACAd) demonstrates a subset of cortical projecting neurons (presumably pyramidal tract projection neurons or PT neurons) generate collateral projections to both the SC.cl and CPc.d. **d)** Circuit schematic illustrating the integration of the cortico-basal ganglia circuit with the cortico-tecto-thalamic circuit. Plus sign circles denote glutamatergic signaling, minus sign circles denote GABAergic signaling. *Abbreviations: CPc.dm, caudoputamen, caudal dorsomedial domain; FG, fluorogold; MOs, secondary motor cortex; MOp, primary motor cortex; LD, laterodorsal nucleus of thalamus; LP, lateroposterior nucleus of thalamus; MD/Hb; mediodorsal nucleus of thalamus/habenula; ORB, orbitofrontal area; PSV, principle sensory nucleus of trigeminal; SNr, substantia nigra pars reticulata; TH, thalamus; SC, superior colliculus; VM-PCN, ventromedial nucleus/pericentral nucleus of thalamus.*

**Supplementary Figure 7.**
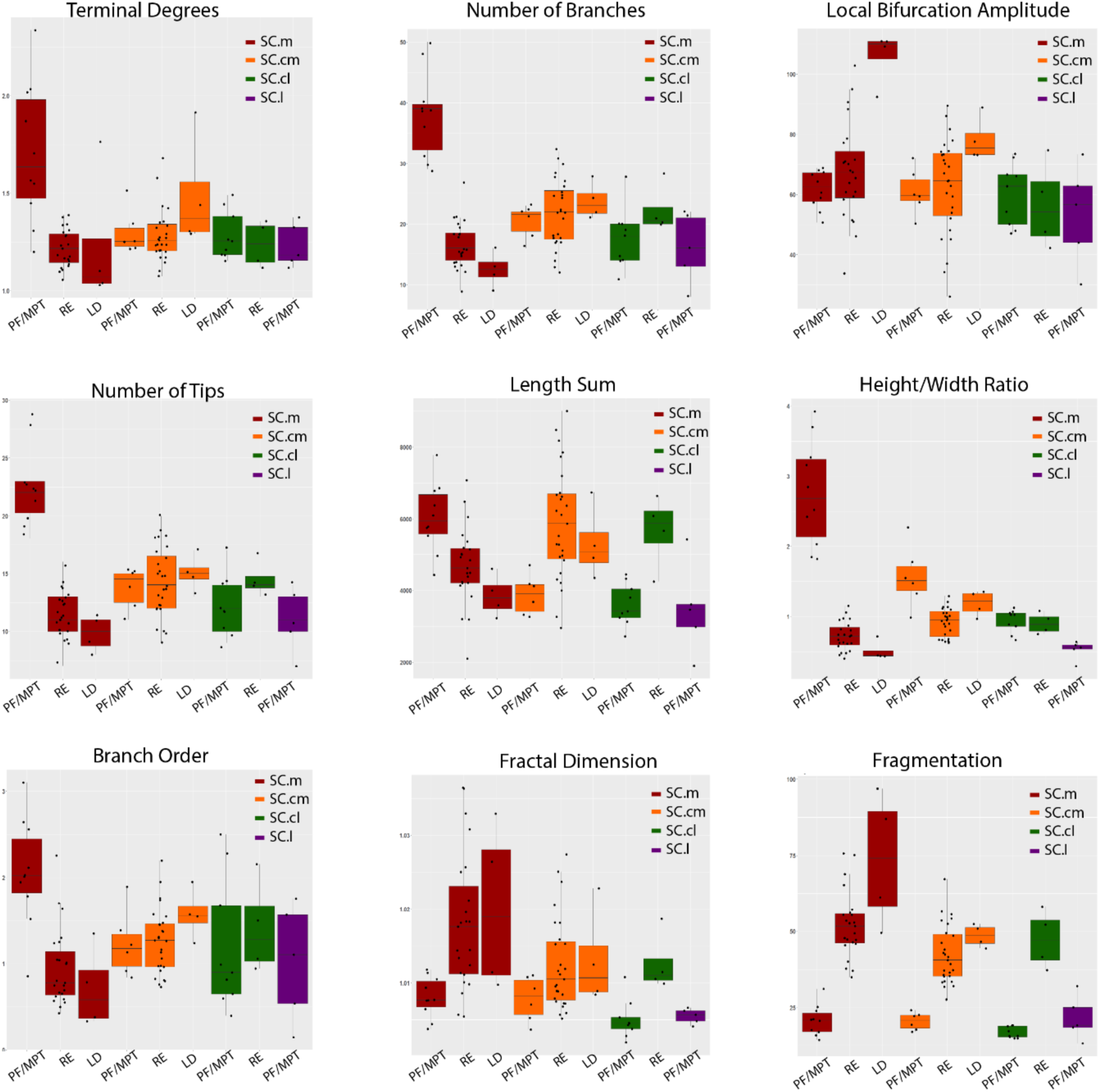
Additional statistical reports from pairwise tests of morphological parameters. All parameters listed survived the false discovery rate (FDR) correction tests. Significant group differences are presented with whisker plots, where the center line represents the median, box limits show the upper and lower quartiles and the whiskers represent the minimum and maximum values. Number SC projection neuron reconstructions to RE (n=54), LD (n=8), and PF/MPT (n=31) (total n= 93 neurons).

**Supplementary Table 1.**
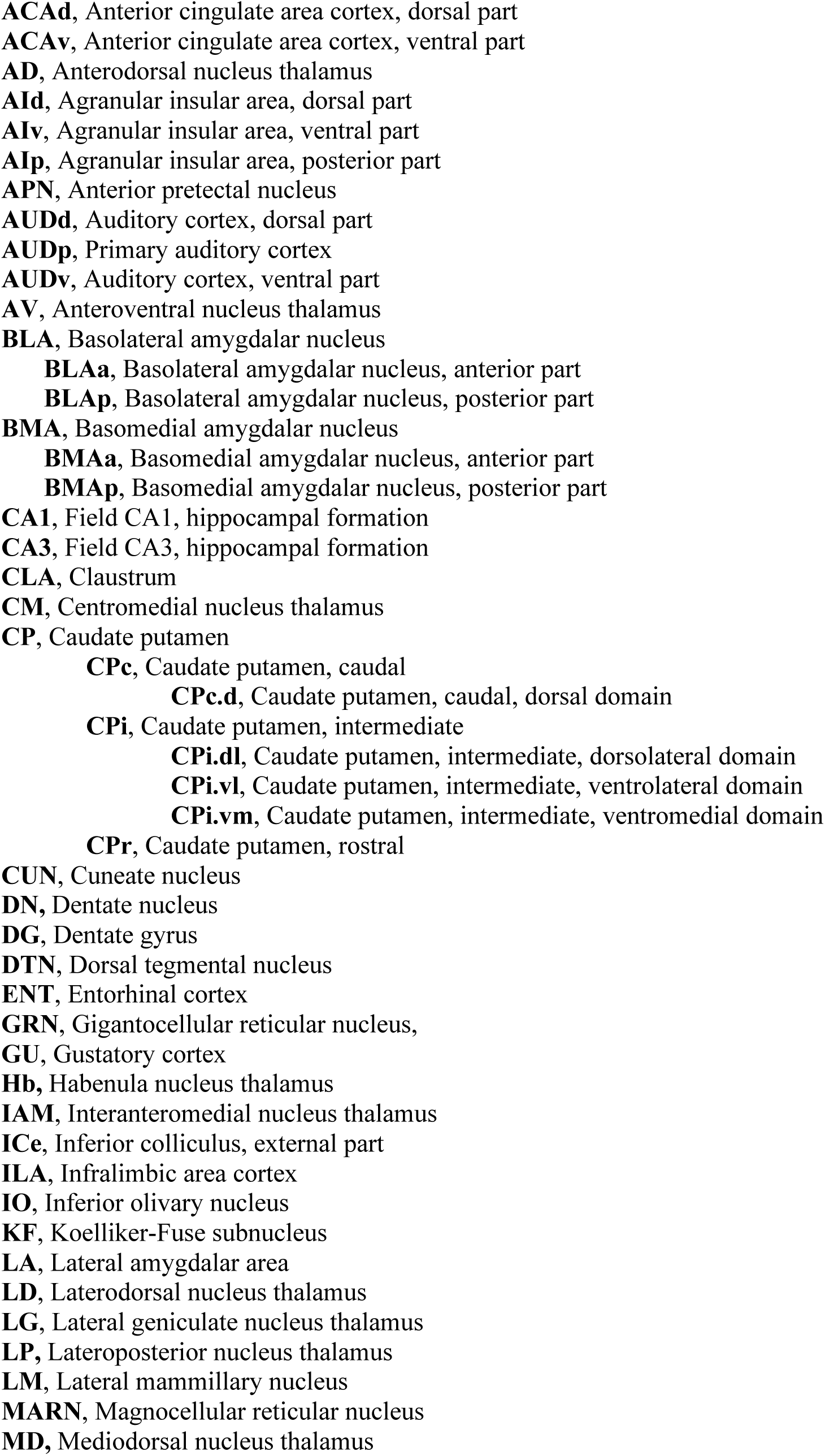
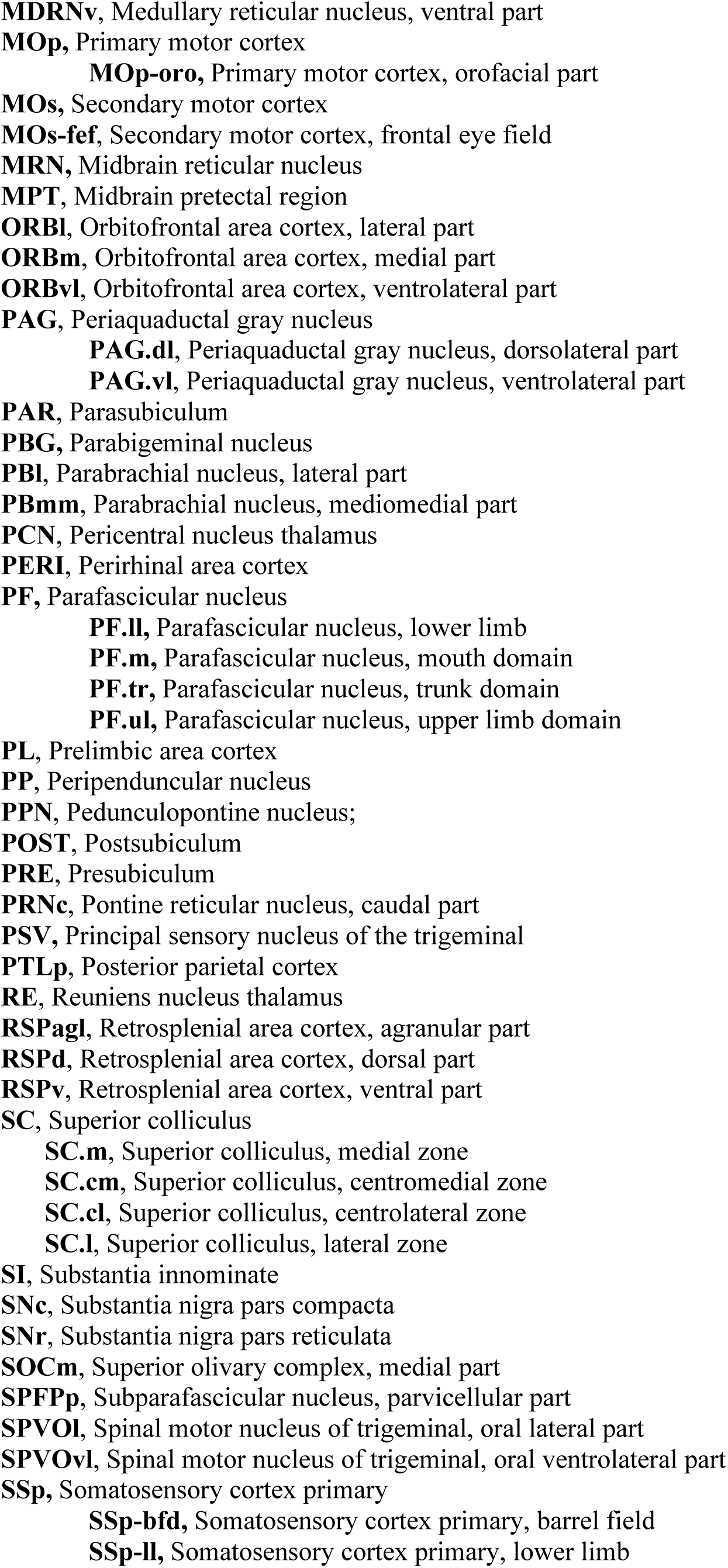
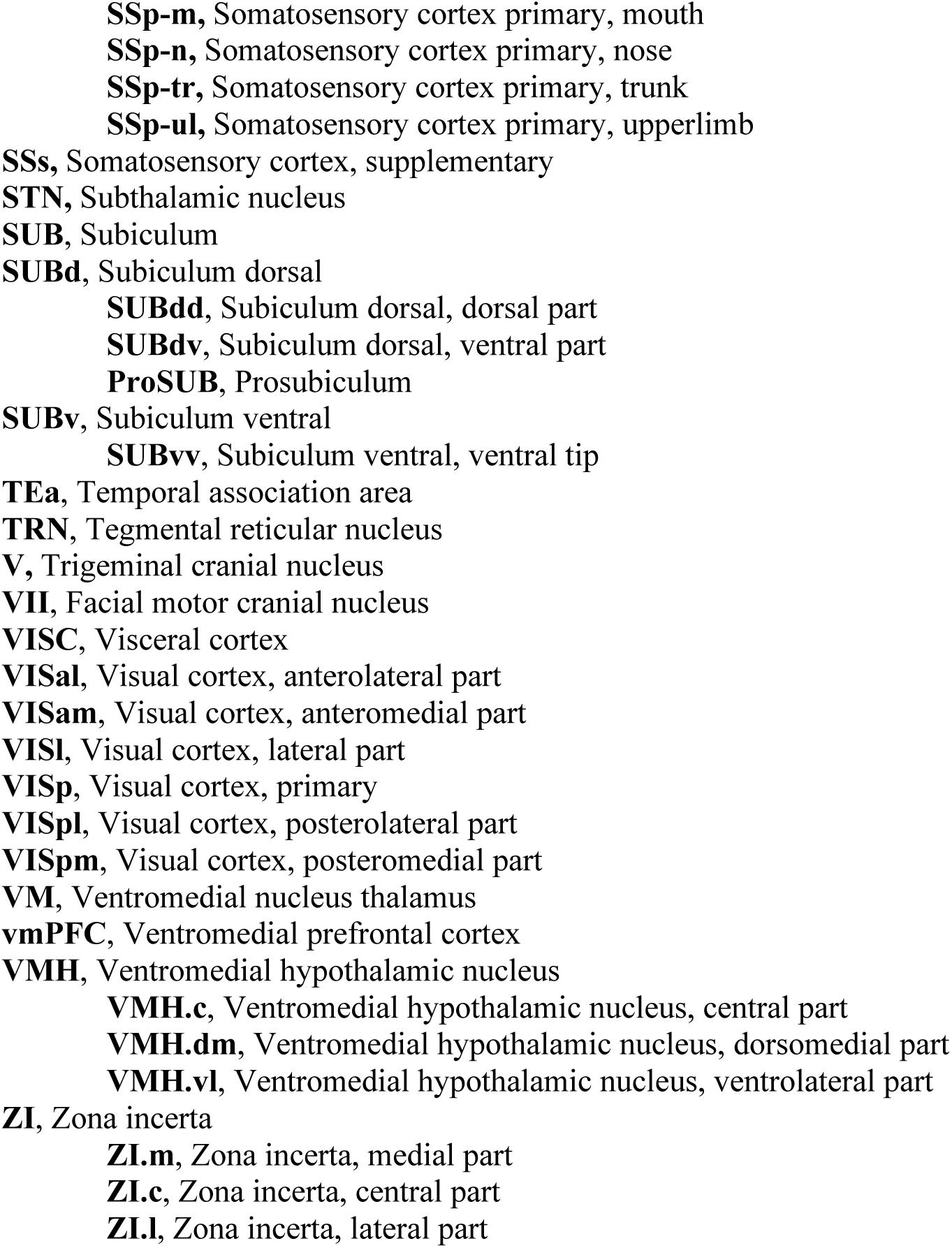
Abbreviation List

**Supplementary Table 2.**
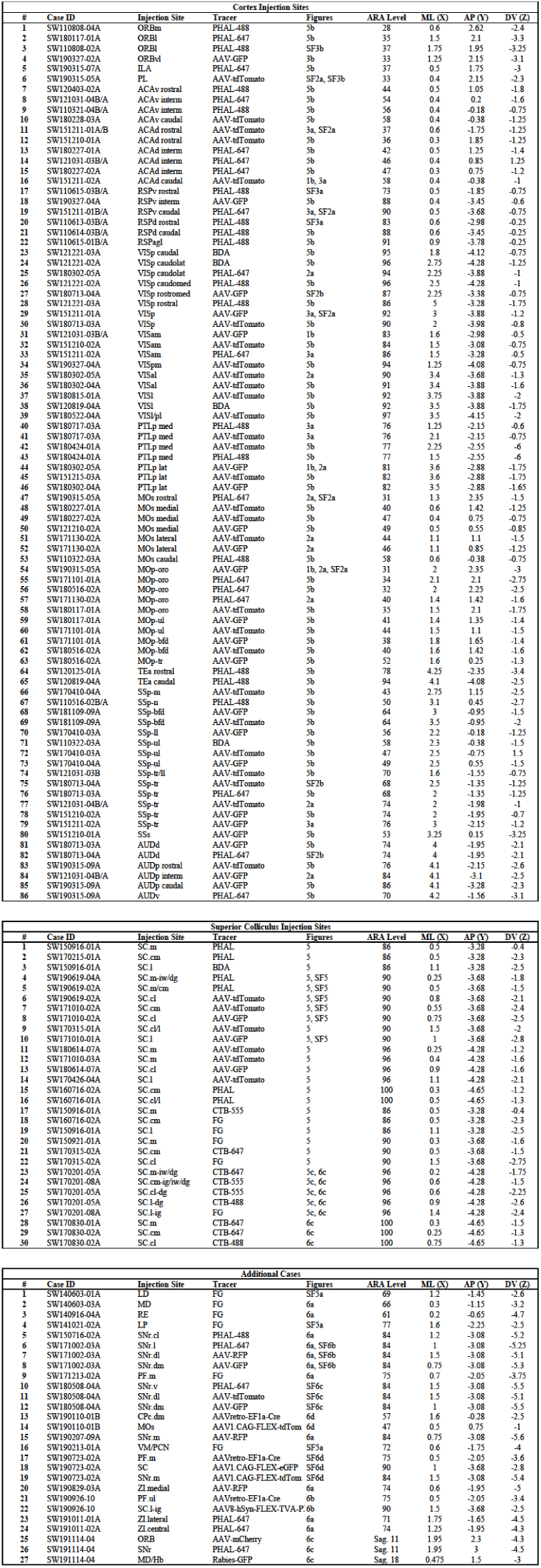
Injection site coordinates and tracer information for cases used throughout the study. Cortex cases can be cross-referenced to flat map injection site location in **Supplementary Figure 1.** Superior colliculus cases can be cross-referenced to injection site map in **Supplementary Figure 4.** (Final formatted Excel file available upon publication.)

**Supplementary Table 3.**
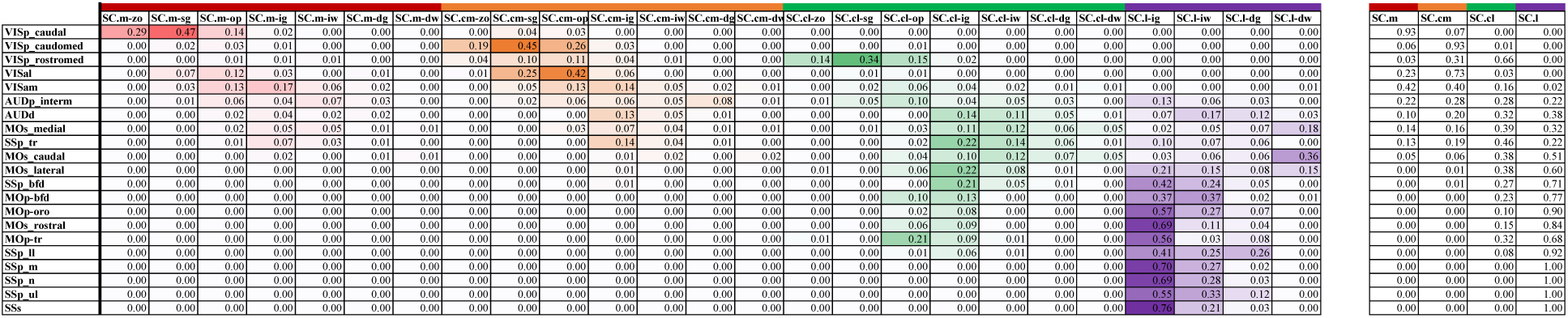
Proportion of labeling values: sensory cortex ROIs to SC zones-layers. Far right summation table corresponds to values used in **Figure 2c**. (Final formatted Excel file available upon publication.)

**Supplementary Table 4.**
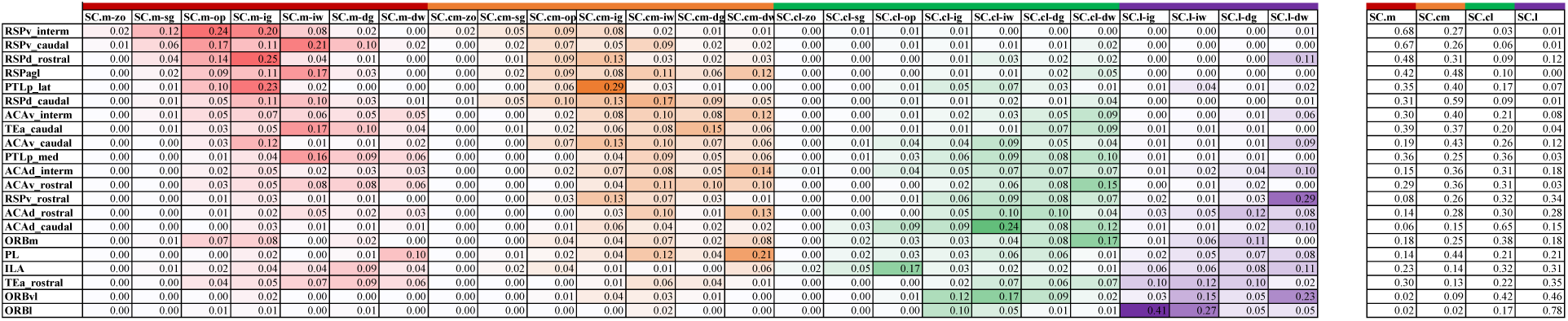
Proportion of labeling values: higher order cortex ROIs to SC zones-layers. Far right summation table corresponds to values used in **Figure 3d**. (Final formatted Excel file available upon publication.)

**Supplementary Table 5.**
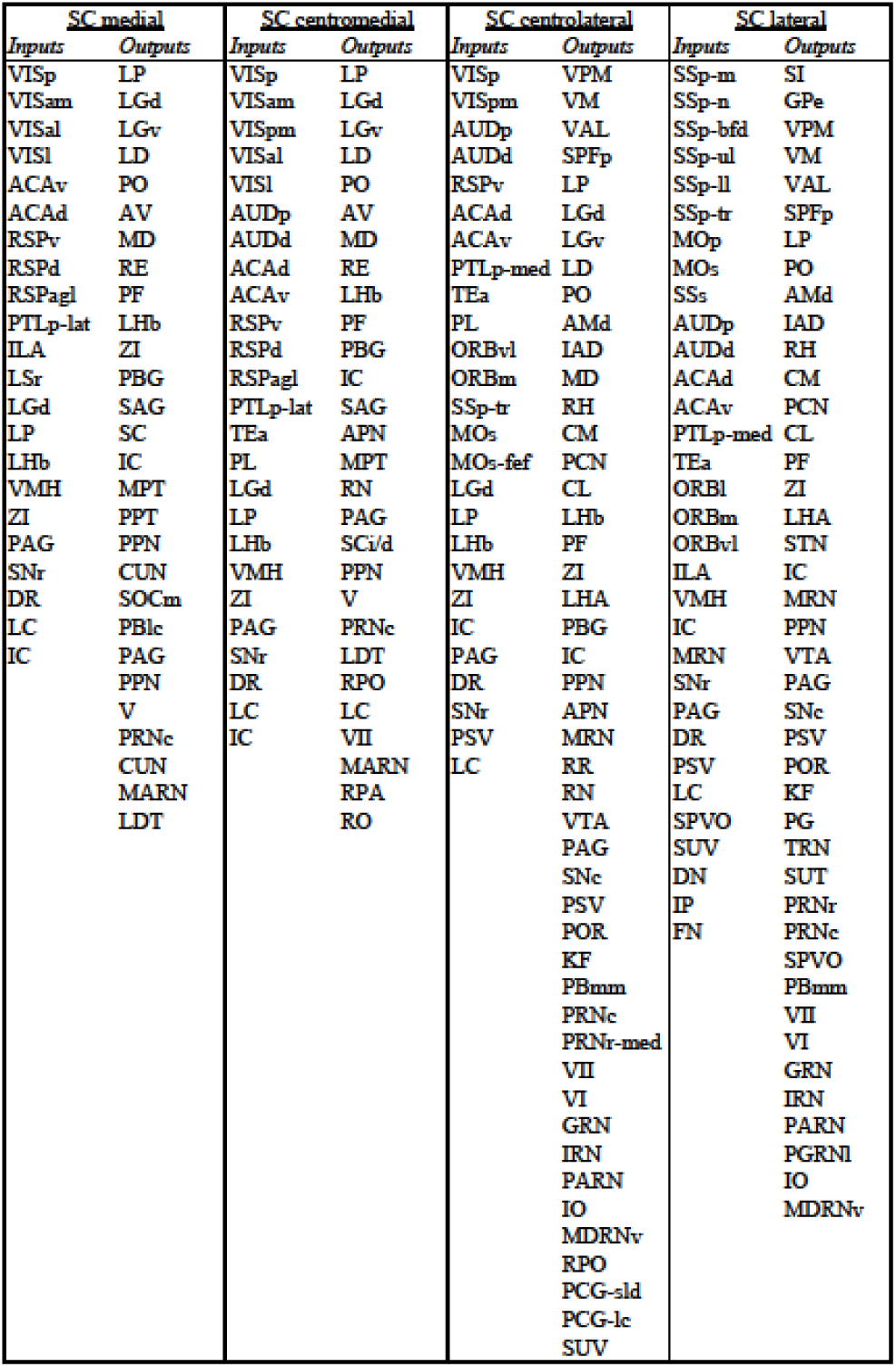
Annotation of input and outputs to each SC zone. (Final formatted Excel file available upon publication.)

**Supplementary Table 6.**
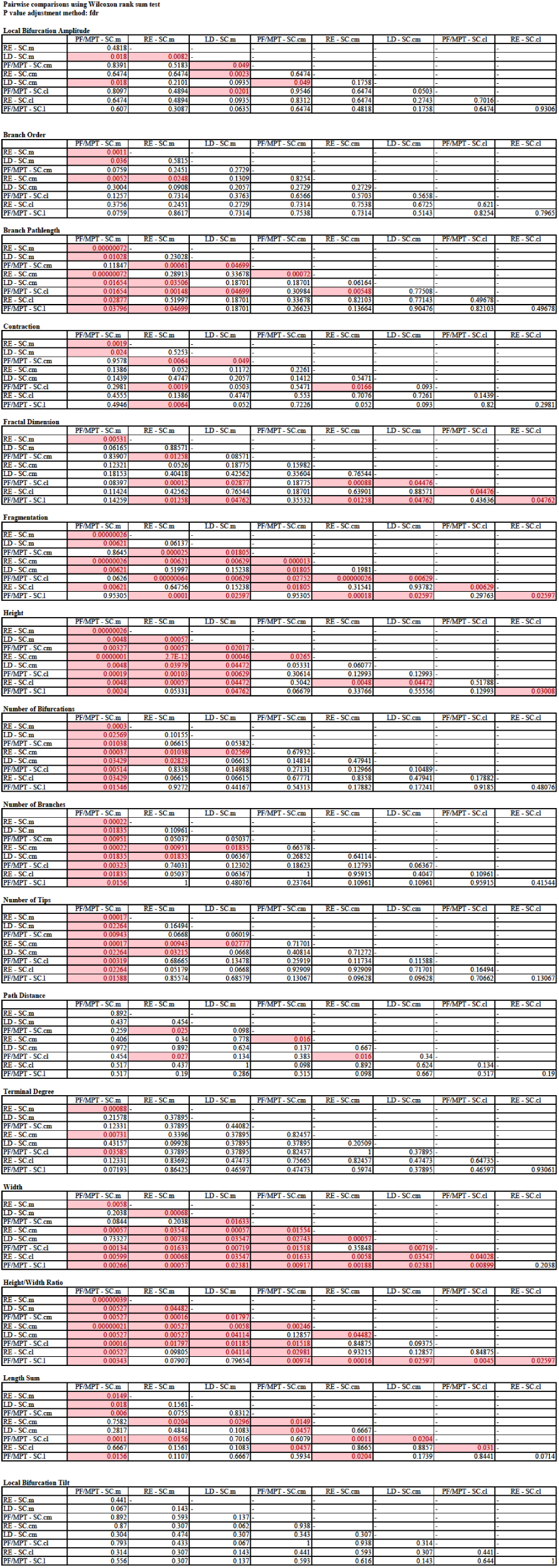
Pairwise p-value table for SC neuron morphometrics. (Final formatted Excel file available upon publication.)

**Supplementary Table 7.**
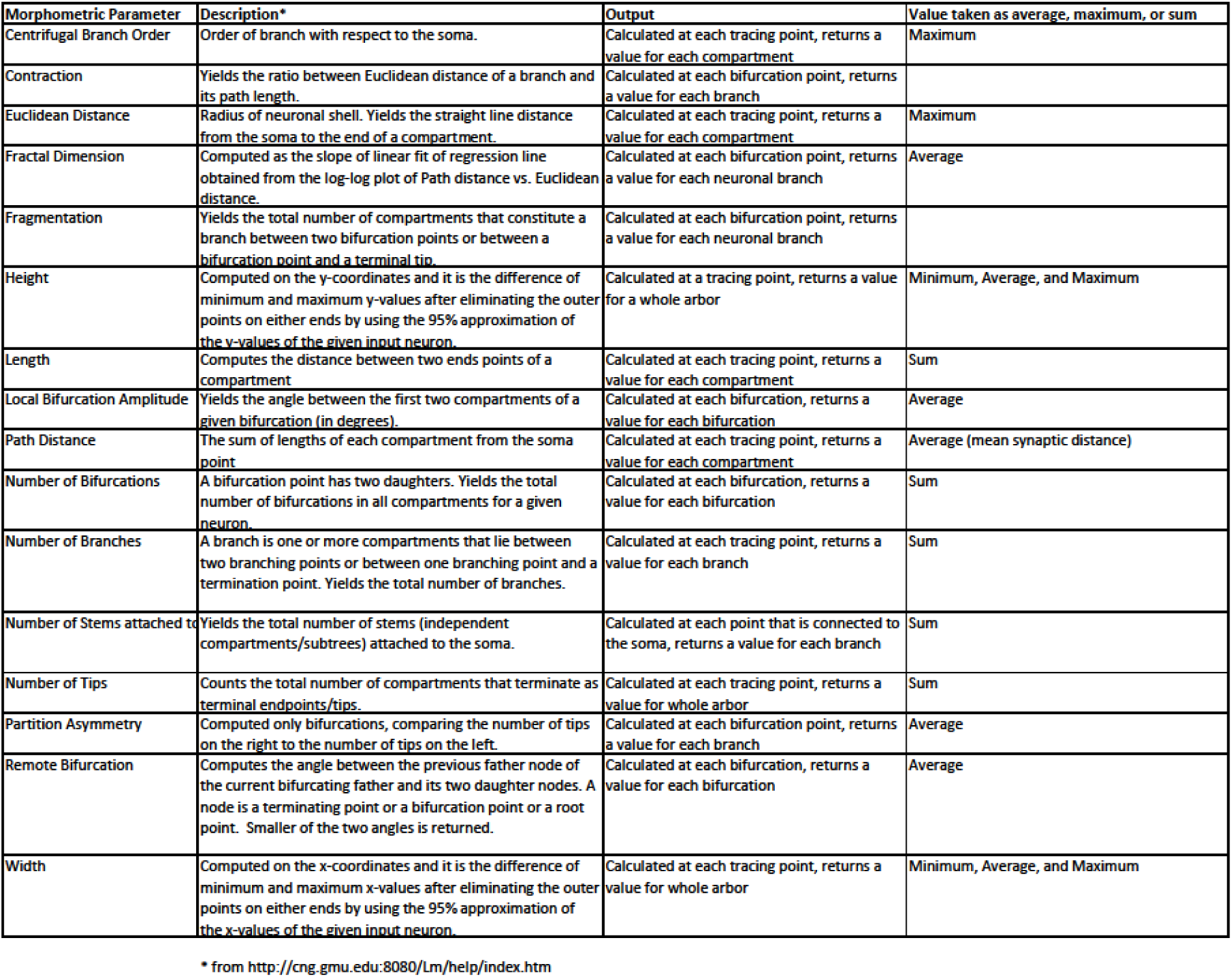
Morphology parameter descriptions, adapted from L-Measure list. (Final formatted Excel file available upon publication.)

## REFERENCES

1. Zingg, B. et al. AAV-Mediated Anterograde Transsynaptic Tagging: Mapping Corticocollicular Input-Defined Neural Pathways for Defense Behaviors. Neuron 93, 33–47 (2017).

2. Zhang, S. et al. Organization of long-range inputs and outputs of frontal cortex for top-down control. Nat. Neurosci. 19, 1733–1742 (2016).

3. Drager, U. C. & Hubel, D. H. Responses to visual stimulation and relationship between visual, auditory, and somatosensory inputs in mouse superior colliculus. J. Neurophysiol. 38, 690–713 (1975).

4. Hadjikhani, N. et al. Look me in the eyes: Constraining gaze in the eye-region provokes abnormally high subcortical activation in autism. Sci. Rep. 7, 1–7 (2017).

5. Klein, C., Raschke, A. & Brandenbusch, A. Development of pro- and antisaccades in children with attention-deficit hyperactivity disorder (ADHD) and healthy controls. Psychophysiology 40, 17–28 (2003).

6. Milner, A. D., Foreman, N. P. & Goodale, M. A. Go-left go-right discrimination performance and distractibility following lesions of prefrontal cortex or superior colliculus in stumptail macaques. Neuropsychologia 16, 381–390 (1978).

7. O’Driscoll, G. A. et al. Executive functions and methylphenidate response in subtypes of attention-deficit/hyperactivity disorder. Biol. Psychiatry 57, 1452–1460 (2005).

8. Overton, P. G. Collicular dysfunction in attention deficit hyperactivity disorder. Med. Hypotheses 70, 1121–1127 (2008).

9. Sesack, S. R., Deutch, A. Y., Roth, R. H. & Bunney, B. S. Topographical organization of the efferent projections of the medial prefrontal cortex in the rat: An anterograde tract-tracing study with Phaseolus vulgaris leucoagglutinin. J. Comp. Neurol. 290, 213–242 (1989).

10. Savier, E. et al. A molecular mechanism for the topographic alignment of convergent neural maps. Elife 6, 1–23 (2017).

11. Ito, S. & Feldheim, D. A. The mouse superior colliculus: An emerging model for studying circuit formation and function. Front. Neural Circuits 12, 1–11 (2018).

12. Sparks, D. L. Translation of sensory signals into commands for control of saccacid eye movements: Role of primate superior colliculus. Physiol. Rev. 66, 118–171 (1986).

13. Krauzlis, R. J., Lovejoy, L. P. & Zénon, A. Superior Colliculus and Visual Spatial Attention. Annu. Rev. Neurosci. 36, 165–182 (2013).

14. Stein, B. E. Organization of the rodent superior colliculus: Some comparisons with other mammals. Behav. Brain Res. 3, 175–188 (1981).

15. Butler, B. E., Chabot, N. & Lomber, S. G. A quantitative comparison of the hemispheric, areal, and laminar origins of sensory and motor cortical projections to the superior colliculus of the cat. J. Comp. Neurol. 524, 2623–2642 (2016).

16. May, P. J. The mammalian superior colliculus: Laminar structure and connections. Prog. Brain Res. 151, 321–378 (2006).

17. Comoli, E. et al. Segregated anatomical input to sub-regions of the rodent superior colliculus associated with approach and defense. Front. Neuroanat. 6, 1–19 (2012).

18. Savage, M. A., McQuade, R. & Thiele, A. Segregated fronto-cortical and midbrain connections in the mouse and their relation to approach and avoidance orienting behaviors. J. Comp. Neurol. 525, 1980–1999 (2017).

19. Dong, H.-W. The Allen reference atlas: A digital color brain atlas of the C57Bl/6J male mouse. John Wiley & Sons Inc., (2008).

20. Blondel, V. D., Guillaume, J. L., Lambiotte, R. & Lefebvre, E. Fast unfolding of communities in large networks. J. Stat. Mech. Theory Exp. 2008, 0–12 (2008).

21. Lancichinetti, A. & Fortunato, S. Consensus clustering in complex networks. Sci. Rep. 2, (2012).

22. Ascoli, G. A., Donohue, D. E. & Halavi, M. NeuroMorpho.Org: A central resource for neuronal morphologies. J. Neurosci. 27, 9247–9251 (2007).

23. Zingg, B. et al. Neural networks of the mouse neocortex. Cell 156, 1096–1111 (2014).

24. Hintiryan, H. et al. The mouse cortico-striatal projectome. Nat. Neurosci. 19, 1100–1114 (2016).

25. Isa, T. & Saito, Y. The direct visuo-motor pathway in mammalian superior colliculus; novel perspective on the interlaminar connection. Neurosci. Res. 41, 107–113 (2001).

26. Tardif, E., Delacuisine, B., Probst, A. & Clarke, S. Intrinsic connectivity of human superior colliculus. Exp. Brain Res. 166, 316–324 (2005).

27. Hall, W. C. & Lee, P. Interlaminar connections of the superior colliculus in the tree shrew. III: The optic layer. Vis. Neurosci. 14, 647–661 (1997).

28. Wang, Q. & Burkhalter, A. Area Map of Mouse Visual Cortex. J. Comp. Neurol. 502, 339–357 (2007).

29. McHaffie, J. G., Stanford, T. R., Stein, B. E., Coizet, V. & Redgrave, P. Subcortical loops through the basal ganglia. Trends Neurosci. 28, 401–407 (2005).

30. Redgrave, P. et al. Interactions between the midbrain superior colliculus and the basal ganglia. Front. Neuroanat. 4, 1–8 (2010).

31. Schwarz, L. A. et al. Viral-genetic tracing of the input-output organization of a central noradrenaline circuit. Nature 524, 88–92 (2015).

32. Widmer, C. G., Morris-Wiman, J. A. & Calhoun, J. C. Development of trigeminal mesencephalic and motor nuclei in relation to masseter muscle innervation in mice. Dev. Brain Res. 108, 1–11 (1998).

33. Canteras, N. S., Simerly, R. B. & Swanson, L. W. Organization of projections from the ventromedial nucleus of the hypothalamus: A Phaseolus vulgaris-Leucoagglutinin study in the rat. J. Comp. Neurol. 348, 41–79 (1994).

34. May, P. J. & Basso, M. A. Connections between the zona incerta and superior colliculus in the monkey and squirrel. Brain Struct. Funct. 223, 371–390 (2018).

35. Falkner, A. L. & Lin, D. Recent advances in understanding the role of the hypothalamic circuit during aggression. Front. Syst. Neurosci. 8, 1–14 (2014).

36. Hoy, J. L., Bishop, H. I. & Niell, C. M. Defined Cell Types in Superior Colliculus Make Distinct Contributions to Prey Capture Behavior in the Mouse. Curr. Biol. 29, 4130-4138.e5 (2019).

37. Li, Y., Wang, D., Ascoli, G. A., Mitra, P. & Wang, Y. Metrics for comparing neuronal tree shapes based on persistent homology. PLoS One 12, 1–24 (2017).

38. Scorcioni, R., Polavaram, S. & Ascoli, G. A. L-Measure: A web-accessible tool for the analysis, comparison and search of digital reconstructions of neuronal morphologies. Nat. Protoc. 3, 866–876 (2008).

39. Meredith, M. A. & Ramoa, A. S. Intrinsic circuitry of the superior colliculus: Pharmacophysiological identification of horizontally oriented inhibitory interneurons. J. Neurophysiol. 79, 1597–1602 (1998).

40. Mooney, R. D., Klein, B. G., Jacquin, M. F. & Rhoades, R. W. Dendrites of deep layer, somatosensory superior collicular neurons extend into the superficial laminae. Brain Res. 324, 361–365 (1984).

41. Saito, Y. & Isa, T. Electrophysiological and Morphological Properties of Neurons in the Rat Superior Colliculus. I. Neurons in the Intermediate Layer. J. Neurophysiol. 82, 754–767 (1999).

42. Corneil, B. D. & Munoz, D. P. Overt responses during covert orienting. Neuron 82, 1230–1243 (2014).

43. Wolf, A. B. et al. An integrative role for the superior colliculus in selecting targets for movements. J. Neurophysiol. 114, 2118–2131 (2015).

44. Stubblefield, E. A., Costabile, J. D. & Felsen, G. Optogenetic investigation of the role of the superior colliculus in orienting movements. Behav. Brain Res. 255, 55–63 (2013).

45. Masullo, L. et al. Genetically Defined Functional Modules for Spatial Orienting in the Mouse Superior Colliculus. Curr. Biol. 29, 2892-2904.e8 (2019).

46. Karabelas, A. B. & Moschovakis, A. K. Nigral inhibitory termination on efferent neurons of the superior colliculus: An intracellular horseradish peroxidase study in the cat. J. Comp. Neurol. 239, 309–329 (1985).

47. Nail-Boucherie, K. et al. Evidence for a role of the parafascicular nucleus of the thalamus in the control of epileptic seizures by the superior colliculus. Epilepsia 46, 141–145 (2005).

48. Shires, J., Joshi, S. & Basso, M. A. Shedding new light on the role of the basal ganglia-superior colliculus pathway in eye movements. Curr. Opin. Neurobiol. 20, 717–725 (2010).

49. Chatterjee, S. et al. Nontoxic, double-deletion-mutant rabies viral vectors for retrograde targeting of projection neurons. Nat. Neurosci. 21, 638–646 (2018).

50. Wickersham, I. R., Sullivan, H. A. & Seung, H. S. Production of glycoprotein-deleted rabies viruses for monosynaptic tracing and high-level gene expression in neurons. Nat. Protoc. 5, 595–606 (2010).

51. Hall, W. C. & May, P. J. The Anatomical Basis for Sensorimotor Transformations in the Superior Colliculus. Contributions to Sensory Physiology Vol. 8 (Academic Press, Inc., 1984).

52. Stein, B. E., Wallace, M. W., Stanford, T. R. & Jiang, W. Cortex governs multisensory integration in the midbrain. Neuroscientist 8, 306–314 (2002).

53. Gale, S. D. & Murphy, G. J. Distinct representation and distribution of visual information by specific cell types in mouse superficial superior colliculus. J. Neurosci. 34, 13458–13471 (2014).

54. Bienkowski, M. S. et al. Integration of gene expression and brain-wide connectivity reveals the multiscale organization of mouse hippocampal networks. Nat. Neurosci. 21, 1628–1643 (2018).

55. Taube, J. S. The Head Direction Signal: Origins and Sensory-Motor Integration. Annu. Rev. Neurosci. 30, 181–207 (2007).

56. Jankowski, M. M. et al. Nucleus reuniens of the thalamus contains head direction cells. Elife 3, 1–10 (2014).

57. Wall, P. M. et al. Differential effects of infralimbic vs. ventromedial orbital PFC lidocaine infusions in CD-1 mice on defensive responding in the mouse defense test battery and rat exposure test. Brain Res. 1020, 73–85 (2004).

58. Illing, R. B. & Graybiel, A. M. Convergence of afferents from frontal cortex and substantia nigra onto acetylcholinesterase-rich patches of the cat’s superior colliculus. Neuroscience 14, (1985).

59. Beninato, M. & Spencer, R. F. A cholinergic projection to the rat superior colliculus demonstrated by retrograde transport of horseradish peroxidase and choline acetyltransferase immunohistochemistry. J. Comp. Neurol. 253, 525–538 (1986).

60. Mátyás, F., Lee, J., Shin, H. S. & Acsády, L. The fear circuit of the mouse forebrain: Connections between the mediodorsal thalamus, frontal cortices and basolateral amygdala. Eur. J. Neurosci. 39, 1810–1823 (2014).

61. Hikosaka, O., Takikawa, Y. & Kawagoe, R. Role of the basal ganglia in the control of purposive saccadic eye movements. Physiol. Rev. 80, 953–978 (2000).

62. Kolling, N., Behrens, T. E. J., Mars, R. B. & Rushworth, M. F. S. Neural mechanisms of foraging. Science (80-.). 335, 95–98 (2012).

63. Petreanu, L., Mao, T., Sternson, S. M. & Svoboda, K. The subcellular organization of neocortical excitatory connections. Nature 457, 1142–1145 (2009).

64. Swanson, L. W. Cerebral hemisphere regulation of motivated behavior. Brain Research (2000).

65. Lin, D. et al. Functional identification of an aggression locus in the mouse hypothalamus. Nature 470, 221–227 (2011).

66. Kunwar, P. S. et al. Ventromedial hypothalamic neurons control a defensive emotion state. Elife 2015, 1–30 (2015).

67. Wei, P. et al. Processing of visually evoked innate fear by a non-canonical thalamic pathway. Nat. Commun. 6, (2015).

68. Shang, C. et al. A parvalbumin-positive excitatory visual pathway to trigger fear responses in mice. Science (80-.). 348, (2015).

69. Van Der Werf, Y. D., Witter, M. P. & Groenewegen, H. J. The intralaminar and midline nuclei of the thalamus. Anatomical and functional evidence for participation in processes of arousal and awareness. Brain Research Reviews vol. 39 (2002).

70. Salay, L. D., Ishiko, N. & Huberman, A. D. A midline thalamic circuit determines reactions to visual threat. Nature 557, (2018).

71. Hintiryan, H. et al. Connectivity characterization of the mouse basolateral amygdalar complex. bioRxiv (2019).

72. Benevento, L. A. & Fallon, J. H. The ascending projections of the superior colliculus in the rhesus monkey (Macaca mulatta). J. Comp. Neurol. 160, 339–361 (1975).

73. Bienkowski, M. S. et al. Extrastriate connectivity of the mouse dorsal lateral geniculate thalamic nucleus. J. Comp. Neurol. 527, 1419–1442 (2019).

74. Kim, S. Y. et al. Stochastic electrotransport selectively enhances the transport of highly electromobile molecules. Proc. Natl. Acad. Sci. U. S. A. 112, E6274–E6283 (2015).

75. Park, Y. G. et al. Protection of tissue physicochemical properties using polyfunctional crosslinkers. Nat. Biotechnol. 37, 73 (2019).

76. Cabeen, R., Laidlaw, D. H. & Toga, A. W. Quantitative imaging toolkit: Software for interactive 3D visualization, data exploration, and computational analysis of neuroimaging datasets. (2018).

77. Feng, L., Zhao, Ti. & Kim, J. neuTube 1.0: A New Design for Efficient Neuron Reconstruction Software Based on the SWC Format. eNeuro (2015).

